# Social survival: humpback whales (*Megaptera novaeangliae*) use social structure to partition ecological niches within proposed critical habitat

**DOI:** 10.1101/2020.12.31.424937

**Authors:** Janie Wray, Eric Keen, Éadin N. O’Mahony

## Abstract

Animal culture and social bonds are relevant to wildlife conservation because they influence patterns of geography, behavior, and strategies of survival. Numerous examples of socially-driven habitat partitioning and ecological-niche specialization can be found among vertebrates, including toothed whales. But such social-ecological dynamics, described here as ‘social niche partitioning’, are not known among baleen whales, whose societies -- particularly on foraging grounds -- are largely perceived as unstructured and incidental to matters of habitat use and conservation. However, through 16 years of behavioral and photo-identification observations of humpback whales (*Megaptera novaeangliae*) feeding within a fjord system in British Columbia, Canada, we have documented long-term pair bonds (lasting up to 12 years) as well as a complex societal structure, which corresponds closely to persistent patterns in feeding strategy, long-term site fidelity (extended seasonal occupancy and annual rate of return up to 75%), specific geographic preferences within the fjord system, and other forms of habitat use. Randomization tests of network congruency and clustering algorithms were used to test for overlap in patterns of social structure and habitat use, which confirmed the occurrence of social niche partitioning on the feeding grounds of this baleen whale. In addition, we document the extensive practice of group bubble net feeding in Pacific Canada. This coordinated feeding behavior was found to strongly mediate the social structure and habitat use within this humpback whale society. Additionally, during our 2004 – 2019 study, we observed a shift in social network structure in 2010 – 2012, which corresponded with environmental and demographic shifts including a sudden decline in the population’s calving rate. Our findings indicate that the social lives of humpback whales, and perhaps baleen whales generally, are more complex than previously supposed and should be a primary consideration in the assessment of potential impacts to important habitat.

## INTRODUCTION

Patterns of habitat use and the factors that determine it are of fundamental interest in animal ecology [1]. As species occupy and use habitats according to various needs, they are establishing a certain ecological niche within a common domain of limited resources [2,3]. To persist in shared space, sympatric species may differentiate modes of habitat use over time to secure resources [4]. Within a single population, individuals may even vary their movements, prey preferences, and behavior in order to reduce niche overlap and minimize intraspecific competition [5]. Such habitat partitioning, on both inter-species and intra-species levels, facilitates the coexistence of functional groups, increases local biodiversity and ecosystem complexity, and uses available resources more efficiently [4].

In any given species, spatiotemporal distribution and ecological niches are the result of negotiations between environmental dynamics and ecological interactions, as well as intrinsic biological factors such as ancestral body plan, reproductive status, and feeding strategy [6–9]. Social relationships are also an important determinant of habitat use in certain species, particularly within the context of innately social behaviors such as mating and parental care. The geography and seasonality of habitat use is often closely related to mating systems and reproductive calendars, e.g., sea turtles [10], marine iguanids [11], many pinnipeds [12], and most seabirds [13]. Even within a single species, mating systems can change in response to resource availability and habitat type (e.g., *Equus africanus* in North America, [14]).

Among the cetaceans, the influence of social-reproductive behavior upon habitat use has been observed in both odontocetes (toothed whales, porpoises, and dolphins) and mysticetes (baleen whales). Examples include nursery groups of dusky dolphins (*Lagenorhynchus obscurus*) who use shallower waters than other social groups in New Zealand [15], sperm whale (*Physeter macrocephalus*) populations whose seasonal geography is partitioned by sex and age [16], baleen whales who practice large-scale movements between winter breeding areas and summer feeding areas [17], and specifically humpback whales (*Megaptera novaeangliae*) who partition breeding habitat according to maternal status and other social factors [18–29]

Outside of a breeding context, such as foraging, the social component of habitat use may not be recognized unless the species travels within stable family groups (e.g., sperm whales and killer whales, *Orcinus orca*) or large pods (e.g., tropical dolphins) [16,30–32]. But even for relatively solitary species, foraging and other non-reproductive behaviors still occur within a social context [33]. Individuals with similar preferences tend to share common spaces and interact regularly with one another, thus adding a social dimension to their patterns of resource use. These interactions can be merely coincidental or actively sought out and coordinated. An example of the latter is when groups of humpback whales gather together and engage in bubble net feeding to capture schooling fish [34,35]. As a complex feeding strategy that requires cooperation and learned behaviors [35], ‘bubble netting’ is a dramatic example of a habitat use strategy with a strong social component. Over time, as socially intensive behaviors such as bubble netting recur, the bonds that form among individuals can serve to reinforce the ecological similarity of an in-group and exacerbate its differences from out-groups [36]. Again, this process of differentiation could happen coincidentally, via simple attraction [37], or actively, via social selectivity and shared learning [38].

It is in this way that the partitioning of habitat and ecological niche within a population can come to fall along social boundaries, and that social and ecological roles can become mutually reinforcing. We shall refer to this mutually reinforcing overlap of social-, habitat-, and ecological-partitioning as *social niche partitioning*. This process has been observed among various vertebrate groups, including terrestrial mammals, e.g., *Hapalemur griseus* in Madagascar [39], marine mammals, e.g., the sympatric communities of killer whales in the Pacific Northwest and Southern Ocean [31,32] and sperm whales in the Mediterranean [40], as well as in other taxa (see next paragraph).

For many such cases, we know that niche partitioning is the direct result of behavioral traits that are socially learned and shared, and thus constitute a form of *culturally-mediated* niche partitioning [31,41–45]. Examples are found among mountain sheep [46], sea otters [47], tool-using apes and monkeys [48–50], and passerine birds that learn about feeding areas, prey sizes, and predator dangers from parents as well flock members of different species [51,52]. Among the cetaceans, examples can be found in the sperm whales of the Galapagos archipelago [16,45,53–56], ecologically and acoustically specialized pods of killer whales [57], bottlenose dolphins (*Tursiops truncatus*) in Moreton Bay, Australia [58], dusky dolphins of New Zealand’s South Island [59], and tool-using dolphins elsewhere [60]. In these examples, patterns in habitat partitioning and ecological role within a shared space correspond closely to the social structure of the population. Sociality informs habitat use, and *vice versa*.

Among the baleen whales, however, social niche partitioning is rare, if it occurs at all. Biologists are aware of culturally inherited site fidelity to certain breeding and feeding regions (e.g., [61–63]; small, interannually stable social groups with unique habitat and prey preferences in areas separate from the remainder of the population (e.g. the Puget Sound feeding group of gray whales, *Eschrichtius robustus*, [64]); as well as instances of cultural transmission of novel feeding techniques throughout humpback whale social networks (e.g., ‘lobtail’ feeding in the western North Atlantic, [65]; ‘trap feeding’ in southern British Columbia, [66]). However, it is not clear the extent to which social structure and other aspects of habitat use align with these culturally transmitted feeding behaviors. Even on breeding grounds (see citations provided above), social habitat partitioning does not necessarily involve ecological differentiation.

Baleen whales practice wide-ranging movements, carry out most of their behaviors and interactions well below the sea surface, and exhibit minimal sexual dimorphism, all of which makes their social lives particularly difficult to study [67,68]. Their social behaviors are generally considered less stable and less consequential than those of odontocetes [69]. This perception has held for the humpback whale, arguably the best studied baleen whale [17,70], which typically forms fluid fission/fusion groups [71–73] in which long-term social bonds have been seen as the exception [70,72,74]. While this has consistently been the case for social behavior in breeding areas [75,76], an alternative view of humpback social life on feeding grounds is now emerging thanks to evidence of long-term social bonds and social network structure in the Gulf of St. Lawrence [77] and southern Gulf of Maine [78].

Understanding the extent to which such social processes mediate habitat use within humpback whale feeding areas, if at all, is of ecological interest in general, but is also needed for the management of important habitat areas. For example, the fact that humpback mother-calf pairs and Dusky dolphin nursery groups exhibit clear preferences for certain habitat features within breeding areas is useful to managers who must discern how best to allocate limited resources [15,28,79]. Likewise, in feeding areas, social cohesion would be of immediate relevance to managers if it were indeed a factor in habitat partitioning, foraging success, or survival [38,79].

Herein we present evidence of social niche partitioning of a proposed critical habitat for humpback whales in northern British Columbia, based upon 16 years of behavioral observations and photo-identification surveys. We also provide the first documentation, to our knowledge, of long-term social bonds and the extensive practice of bubble net feeding among humpback whales in Pacific Canada, and discuss the implications of these findings for ecology and conservation within those waters.

### Objectives

Our first objective in this study was to describe humpback whale habitat use within a mainland fjord system proposed as critical habitat [80]. Hereafter we use the term ‘habitat use’ to refer collectively to characteristics of both site fidelity and behavior. By ‘site fidelity’, we refer to the return of individuals to the study area across years (interannual fidelity) as well as to the duration of occupancy within any given year. With the term ‘behavior’, we refer to patterns of discrete actions of individuals or small groups (e.g., traveling, resting, bubble net feeding, bringing calves to the area, etc.), to patterns of spatial aggregation associated with those actions, and to geographic and seasonal patterns in the prevalence of those actions.

Second, with the same field effort, we sought to characterize the humpback whale society within this fjord system by answering the following questions: how prevalent are strong social relationships in the local population, and how stable are these relationships within seasons and across years? Furthermore, is there internal structure to the social community, and is this structure stable across years?

Third, we used these data on whale habitat use and sociality to determine the extent to which these aspects of humpback whale life mediate one another within this habitat. To do so, we asked whether aspects of the sociality and habitat use of individuals were strongly related, and if so, in what specific ways. Finally, to test for social niche partitioning, we asked whether population-level patterns in habitat use correspond to the internal structure and interannual stability of the social network. Throughout these analyses, we paid particular attention to group bubble net feeding (an innately social behavior), its relationship to other aspects of habitat use, and its influence upon the social dynamics of the local population.

## METHODS

### Sampling procedure

Data on humpback whale relationships were collected from 2004 to 2019 (n=16 years) within the marine territories of the Gitga’at and Kitasoo/Xai’xais and Haisla First Nations in the Kitimat Fjord System (KFS), mainland British Columbia, Canada (N 52.8 – 53.5, W 129.6 – 128.5; Fig 1). Whales were observed from shore-based and vessel-based platforms. The vessel-based survey methods we used in all years of the study are detailed in Ashe *et al*. [81]. Briefly, preplanned survey routes were conducted using a 7 m skiff as weather permitted from April to November (with occasional trips in February, March, and December) (Fig 1). When humpback whales were detected, groups were approached with caution, all individuals were counted, location and behavior noted, and identification photographs of the underside of their tail flukes were collected with standard DSLR cameras and telephoto lenses, following established protocols for this species (e.g., [82]). Fluke photograph capture was non-systematic in order to maximize the number of whales identified. Shore-based observations occurred at Cetacea Lab (53° 6’17.31”N, 129°11’37.72”W) from 2010 to 2016 the Wall Islets (52°51’29.10”N, 129°20’28.53” W) from 2013 to 2015, and Fin Island Research Station (N 53°13’18.94”N, 129°22’34.77”W) from 2017 to 2019. From these stations, groups were observed if they came close, and identification photographs were taken and catalogued.

**Figure 1.**
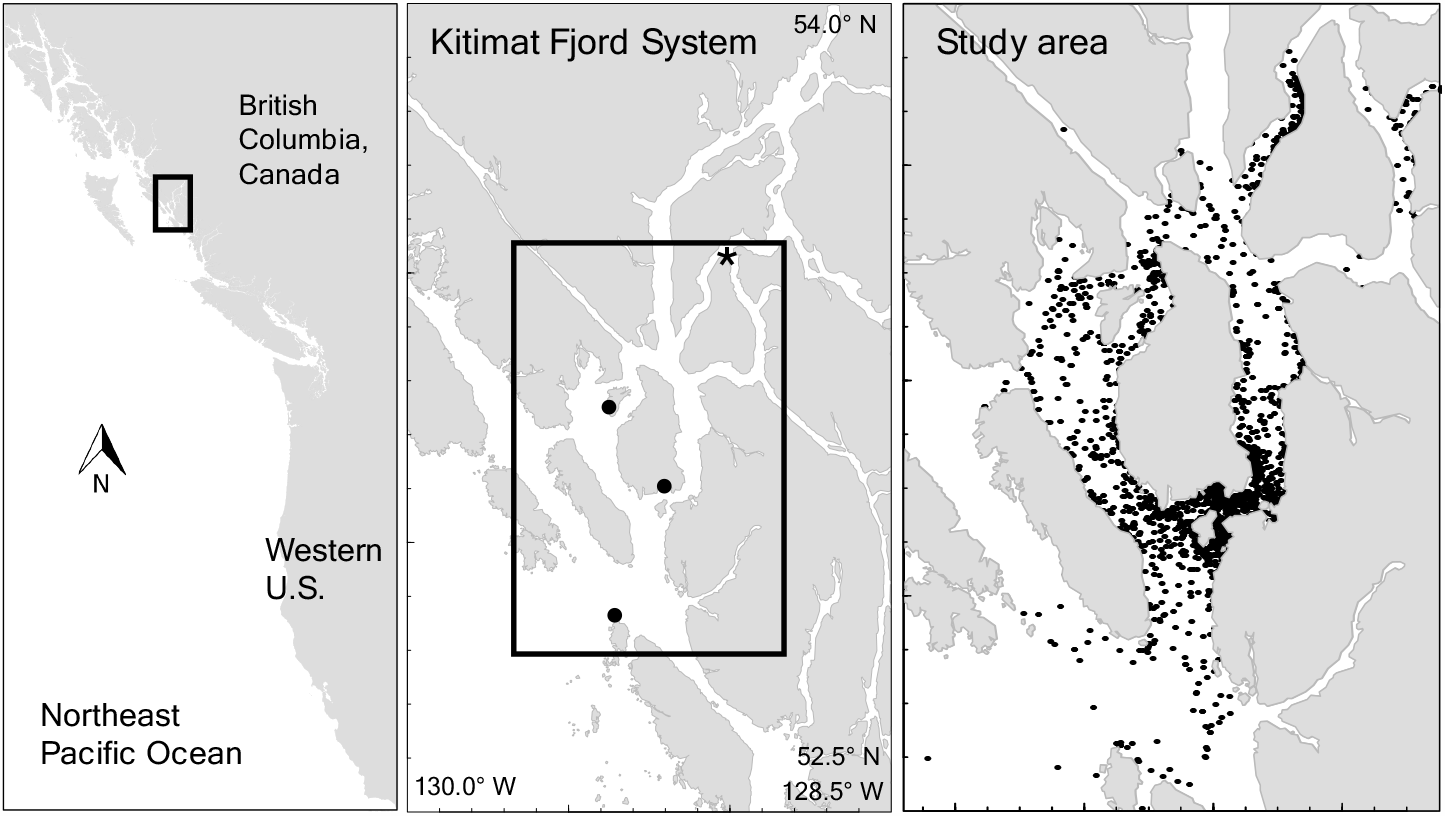
Study area in the Kitimat Fjord System, British Columbia, Canada, within the territories of the Gitga’at and Kitasoo First Nations. *Center pane:* Dots indicate locations of research stations. North to south: Fin Island, Whale Point on Gil Island, and the Wall Islets. Asterisk indicates reference point for fjord position metrics (see main text). *Right pane:* Dots indicate locations of humpback whale photo-identification events, 2004-2019.

### Data preparation

For the purposes of the present analysis, we define a sampling occasion as a single calendar day of photo-identification effort, which usually consisted of up to 16 hours of active monitoring at shore-based stations and 2 – 10 hours of surveying during vessel-based effort. An encounter is defined as a unique observation of a unique group (i.e., if a group of the same social composition was observed twice in one day, it is only included once; if certain whales from a previous group are seen with other whales in a separate group later in the same day, both encounters are included). Females were identified when they arrived with a calf, such that sex is known only for documented mothers in this study. All photographs were scored for quality and were used to populate annual catalogs of identified individuals (after Ashe *et al*. [81]).

For our analysis, sighting histories were developed for each individual that summarized occupancy patterns (see below), observed feeding behaviors, and if a calf was present. To summarize the geographic occurrence of individuals within the KFS, we found their average (and standard deviation) ‘fjord position’, i.e., the mean swimming distance, in km, from an individual to the inland-most point of the study area (53°33’12.11”N, 128°59’55.39”W, which corresponds to the intersection of Verney Passage and Devastation Channel, denoted by the asterisk on Fig. 1). Swimming distance describes the shortest possible route around islands between two points within the KFS (R package ‘bangarang’, [83]; this metric has been used in Keen *et al*. [84] and Keen *et al*. [85]). Small values for average fjord position correspond to whales that are commonly seen deep within the fjord system, whereas large values correspond to whales commonly seen in the outer channels near the continental shelf. These and subsequent analyses, except where noted, were carried out in R 4.0.2 [86].

We defined a group as individuals that come within two body lengths of each other, and coordinating their swimming, diving, and/or ventilation behavior for at least one surfacing, following previous studies [73,77,78,87–90].

During close observation of whale groups, we identified six primary behaviors: bubble net feeding (BNF) was identified by rings of bubbles and feeding at or just below the surface. Milling-feeding was inferred according to several factors: travel pattern was circuitous or repetitively back-and-forth within the same 1 km2; dives were long (more than 5 minutes); and surface sequences comprised many breaths during which the animal was uncommonly still at the surface, suggesting recovery from feeding activity at depth. Traveling was indicated by swimming in one direction for a period of time greater than 30 minutes (or 10 minutes when observed from shore) in which fluking was rare. Resting was indicated by directed travel at low speed, low breathing rate, lingering at or just below the surface, and rare fluking. Sleeping whales were motionless floating at the surface, breathing only one to two times per minute. Robust behaviors included breaching, pectoral or tail slaps, and other energetic surface displays. We used ‘social’ as a designation to capture behaviors during group interactions that were not clearly tied to the other behaviors and were clearly directed towards others in the group who were in close proximity. These included lingering at the surface in very close proximity to each other, body movement or gestures between whales, tonal blows, side rolling, and submerging or surfacing in synchrony. Other logged behaviors included rolling in kelp and interactions with sea lions.

### Habitat use

#### Site fidelity

We characterized population site fidelity based upon the tendency of humpback whales to occupy the study area or to return to it over some period of time [91]. To do so, we used population-level and individual-based metrics of within-season and interannual observations (see Supplementary Appendix 1 for complete details of the analyses in this section).

Population-level residency patterns within a year were examined using Lagged Identification Rates (LIR; [92,93]), which depict the probability that an individual identified on any given day will be re-identified *τ* = 1, 2, 3, …, *τ*_max_ days hence. We set *τ*_max_ to 230 days within the same year as *τ* = 0 (chosen because the maximum duration of any field season was 223 days), used only lags with 10 or more paired identifications to build the LIR curve, and obtained confidence intervals using 100 bootstrap replicates of the data [94]. We used permutation tests of the data stream [95] to evaluate the statistical significance of the observed LIR curve using a null model of random movement in and out of the study area. For this and all subsequent null model tests, we confirmed that our randomization set size was sufficient to stabilize the p-value. To identify the most likely factors affecting LIR dynamics, we used SOCPROG 2.9 [96] to fit exponential decay curves to observed LIRs using a combination of modeled demographic parameters, including population size, mean residence time, and rates of emigration, immigration, and mortality (after Whitehead [97], Whitehead [94], Ramp *et al*. [77], and Perryman *et al*. [93]).

Individual patterns in seasonal occupancy were characterized using standard indicators of site fidelity, including occurrence, permanence, and periodicity as defined in Tschopp *et al*. [98], as well as the Standardized Site Fidelity Index (SSFI) developed by Tschopp *et al*. [98]. To characterize interannual site fidelity, we calculated the population’s annual return rate (number of recaptures of whales seen in previous years divided by the total number of captures in the current year) for each year of the study (*sensu* Acevedo *et al*. [99]), and determined the proportion of study years in which each individual was seen.

We applied a Kruskall-Wallis rank sum test to ask whether site fidelity metrics for known bubble net feeders differed from the remainder of the identified population (i.e. those we never observed bubble net feeding). We then used permutation tests to determine whether the LIR of bubble net feeders differed with statistical significance from that of the remainder of the identified population.

### Behavior

We characterized the geography and clustering patterns of discrete behaviors using a combination of permutation tests and Generalized Linear Mixed Models (Supplementary Appendix 2). To test for patterns in the geography of behavior execution, encounters were pooled according to their average fjord position in bins of 10 km, from 0 km to 100 km. For each variable of interest (e.g., bubble-net feeding), the proportion of encounters in which whales exhibited this behavior (‘behavior rate’) was noted within each distance bin. We then used permutation tests to determine whether this rate was statistically significant. We carried out this test for the following variables: bubble net feeding, collectively all other modes of feeding, social activity / posturing, resting / sleeping, the presence of a known mother, the presence of a calf, average group size and calendar day of encounters. To test whether observed differences in (minimum) cluster size among behavior categories are statistically significant, we built a Poisson Generalized Linear Mixed Model (GLMM) in which behavior categories were included as a fixed effect and whale identification and study year were included as random effects.

### Sociality

The prevalence and stability of social relationships were characterized using weighted association indices and Lagged Association Rates (LARs). We selected the Simple Ratio association Index (SRI, [100]) because 1) we lacked calibration data and 2) the biases inherent to the SRI are more predictable than to an alternative such as the Half-Weight Index [101,102].

#### Stability of associations

We calculated LARs using SOCPROG 2.9 and the R package “asnipe” [103] to describe the temporal stability of relationships over time ([92]; Supplementary Appendix 3). For these analyses, we subset our catalog to those individuals seen 10 or more times and used the same sampling periods and maximum time lags as in the LIR analysis above. We tested whether dyadic stability differs between whales that practice bubble net feeding and those that we have not observed doing so. This test involved calculating several LAR curves: one for all groups encountered, a second for groups that contained at least one known bubble net feeder (these groups could also contain whales not known to bubble net), and a third for groups that contained whales not known to bubble net feed. As with the LIR analysis above, we used permutation tests to evaluate the significance of LAR observations and conventional decay model fitting in SOCPROG 2.9 to identify the feasible social processes underlying the temporal stability of associations.

#### Social differentiation

We used maximum likelihood approximation in SOCPROG 2.9 to calculate social differentiation (S): the heterogeneity of social associations described using the variability of the “true” SRIs estimated by a Poisson model [94]. Values of S close to 0 indicate homogenous relationships within the population, values above 0.5 reflect well differentiated social networks, and values greater than 1 indicate strong differentiation. The correlation coefficient, *r*, between S and the observed (measured) AIs was used to determine AI accuracy and their power in testing for social relationships [94].

#### Social preferences

To disentangle social affinities from associations that may not be driven by social preferences, we used generalized affiliation indices (GAIs) that control for non-social factors when constructing network weights ([104]; Supplementary Appendix 4). Predictor variables used in the calculation of GAIs were the following: joint pairwise gregariousness of dyads (after Godde *et al*. [105], with the correction from Whitehead and James [104]); geographic overlap, defined as the proportion of years in which both individuals were identified that they occurred within 15 km of the same fjord position (reflecting the typical longest dimension of a single channel within the KFS); and temporal overlap, defined as the proportion of years in which both individuals were identified, out of the total number of years in which at least one of the two was identified.

#### Significance tests

We used permutation tests to determine the significance of observed indices of association (SRI) and affiliation (GAI) for each dyadic association and across the population of whales identified on at least 5 occasions. The null models from these permutations were used to test the hypotheses that there were strong associations and that preferences were more prevalent and stronger than expected by random chance. We conducted these permutation tests for the following subsets of the humpback whale social network: BNF:BNF (ties among known bubble net feeders); Other:Other (ties among other whales); BNF:Other (ties between known bubble net feeders and other whales).

#### Network structure & stability

We used significant dyadic associations and affiliations to build social network visualizations in the R package ‘igraph’ [106]. To determine the structure of the social network (Supplementary Appendix 5), we selected the Louvain clustering algorithm [107] in ‘igraph’ to allow for multi-level (nested) and/or overlapping communities, then used permutation tests to determine whether the degree of community clustering observed is greater than would be expected from random chance.

To examine the inter-annual stability of network structure, we built association networks for a running four-year interval (2004-2007, 2005-2008, …, 2016-2019; n=12) for the population of humpback whales seen on at least five occasions throughout the 16-year study. We then used permutation tests to compare structural changes to the network during this time series to the range of changes that could be expected based upon random chance.

### Relationship between sociality and habitat use

To determine the extent to which patterns in habitat use were related to the humpback whale social network, we developed separate tests for dyadic associations, the full network of associations, and the full network of social preferences.

#### Dyadic associations

To test whether dyadic associations were behavior-specific, and whether engaging together in a certain behavior increases the chances of engaging in others, we developed the following randomization routine. For all dyads seen together on at least three occasions, we first determined the proportion of occasions (‘behavioral rates’) in which bubble net feeding, other modes of feeding, traveling, resting, and robust behavior were observed. We then took a behavior of interest, e.g., bubble net feeding, and identified the dyads who had been encountered while engaged in that behavior (‘practitioners’) and the dyads that had not (‘others’). Between these two groups, we then compared the rate of a secondary behavior, e.g., traveling, using a bootstrap differencing technique [108] in which random samples were drawn from the two groups and the difference of those samples was recorded, and this process was repeated 10,000 times to produce a distribution of differences. The mean of this distribution was used as the test statistic. We compared it to a null distribution of means produced by randomizing the behavior notes in the original sighting records and re-running the procedure 1,000 times. By comparing the observed mean difference to the null distribution, we determined the probability that dyads engaged in the primary behavior would ever be observed on a separate occasion to be engaged in the secondary behavior. If the observed mean difference fell *above* the 97.5% quantile of the null distribution, practitioners of the behavior of interest (in our example, bubble net feeding) were significantly more likely to be engaged in the secondary behavior (traveling) than would be expected from random chance. If the observed mean difference fell *below* the 2.5% quantile of the null distribution, practitioners were significantly less likely to be engaged in the secondary behavior.

#### Association network

Our test for social niche partitioning within the association network was based upon the concept of network congruency. Based on the clustering algorithm described above (see *Network structure and stability*), we noted the community to which each individual was assigned. Then, for each possible dyad pair, we noted whether or not the two individuals were assigned to the same community.

We then created a null congruency model based on a routine of 1,000 iterations, in which community assignments were shuffled, community membership (same or not) was noted for each dyad pair, and the randomized result was compared to the observed. For each dyad, if individuals either belonged to the same community in both the observed and randomized clusterings *or* belonged to different communities in both clusterings, then that dyad is considered to be congruent across the two networks [109]. The proportion of congruent dyads in the networks is a measure of the agreement, or ‘congruency’, between two clustering schemes. We carried out this congruency comparison using the igraph function ‘compare’ (method = “adjusted.rand”; [106]). In this way, the randomizations produced a congruency distribution under the null hypothesis of random network clustering, in which positive values indicate better congruency than would be expected by random chance [110].

We then used k-means clustering to assign individuals to communities based upon standardized numerical variables that characterize some behavioral components of habitat use: mean and standard deviation of fjord position (i.e., distance from the inner fjord), bubble net feeding rate (the proportion of encounters in which the individual was engaged in bubble net feeding), feeding rate (referring to modes of feeding other than bubble netting), social rate, and resting rate. We considered other variables (e.g., observed calving rate and average minimum group size) but excluded them based on their collinearity with more informative variables (e.g., observed calving rate is a function of times seen and therefore site fidelity; see next stage of analysis. Also, group size is correlated to certain behaviors such as bubble net feeding, Table S1). For every possible combination of these variables (n=63), we used k-means clustering in R (base ‘stats’ package; maximum iterations = 100, number of starts = 10) to assign each individual into *k*=7 communities, which is the number of communities identified in the association network structure analysis described above (see *Results*).

We then calculated the congruency of these 63 behavior-based clusterings with the original social clustering scheme. For each, the congruency index was then compared to the null congruency distribution to determine its significance: the proportion of randomized congruency indices that was greater than the behavioral congruency index was treated as a p-value. The significance (α=0.05) of a k-means clustering scheme, which is based upon a certain subset of behavioral variables, indicates that exhibition of those behaviors across the population is, in fact, correlated to its social structure.

If several k-means clustering schemes were significant, we evaluated them based upon their ranked decline in congruency performance (Δ Adjusted Rand Index; [111]). The variables included in this significance set were treated as important behavioral correlates of social network structure, and the proportion of significant clusterings in which each variable was included was treated as a rough qualitative measure of variable importance.

To double-check this social-behavioral congruency significance test, we conducted the process in reverse. We built a randomization set of k-means clustering schemes (n=1,000) by shuffling each behavior variable with respect to humpback whale ID (we did this only for the variable set that yielded the highest congruence metric of the 63 combinations tested), then reassigning individuals to one of *k=*7 communities using the k-means algorithm. We compared these randomized clusterings to the social clustering using the Adjusted Rand Index, then compared this null distribution to the realized social-behavioral congruency. If less than 5% of the null congruency distribution was greater than the realized value, the correlation of social and behavioral clusterings was validated.

This k-means clustering and significance testing process was repeated for site fidelity variables (proportion of years seen, mean arrival date, mean minimum stay, and Standardized Site Fidelity Index; n=15 variable combinations), which were also standardized prior to clustering. Finally, the behavior and site fidelity variable sets were combined and k-means clustering was repeated to determine which variable combination produced the highest congruency with the social network.

#### Preference network

Since the Generalized Affiliation Index (GAI) is designed to isolate social preferences from the effects of temporal and geographic overlap in the study area, cluster congruency tests may not be the appropriate approach for testing the interaction between habitat use variables and social relationships. Instead we used two analytical approaches: first, assortativity coefficient (AC) significance testing, and second, network position ∼ trait correlation tests (after Farine [95] and Perryman *et al*. [93]).

##### Assortativity coefficients

We asked whether the GAI network was assorted according to tendencies in behavior and/or site fidelity. To do so, we calculated ACs for each candidate variable using R package ‘assortnet’ [112]. ACs are positive if individuals with similar traits tend to positively connect, and negative if strongly different individuals tend to negatively connect (i.e., avoid each other). Since both GAIs and ACs can be positive or negative, we conducted this assortativity analysis separately for the network of social affiliations (positive GAIs) and the network of social avoidance (negative GAIs).

The significance of each observed AC was determined by comparing it to a null distribution of ACs that would be expected if assortment by a given trait with the network was merely random (1,000 iterations, based upon node shuffling; [93,113]). Within this framework, in the case of positive GAIs, ACs that are significantly larger than expected serve to indicate that trait similarity relates closely to social preferences within the society. In the case of negative GAIs, ACs that are significantly *more extreme* than expected indicate that dissimilar individuals strongly avoid each other, while ACs that are significantly *less extreme* than expected indicate that similarity between individuals reduces avoidance and leads to assortment.

##### Network positions

To investigate whether individual social patterns are related to strategies of habitat use, we calculated position metrics for individuals within the affiliation network. We used the ‘tnet’ package [114] in R to calculate each individual’s weighted degree (summed weight of all connections), weighted betweenness centrality (measure of how often an individual is located on the shortest path between two others, favoring short paths of weaker ties rather than longer paths of stronger ties; α tuning parameter = 0.5), closeness centrality (the degrees of social separation between one whale and all other individuals), and local clustering coefficient (measure of neighborhood completeness). We treated all negative GAIs as zeroes since we were primarily interested in the effect of positive social preferences.

If social and habitat-use strategies are prevalently interrelated in common ways, then individuals with certain traits may, on average, share similar network positions. We tested this hypothesis with respect to several trait variables by comparing the slope coefficients of linear models built with the observed data (Observed network position metric ∼ Observed trait) and with randomized trait data (Observed network position metric ∼ Randomized trait; n=1,000; after Farine [95]). Observed slopes that fell outside of the 0.025 – 0.975 quantiles of the randomized sets were considered significant.

## RESULTS

### Data summary

A total of 5,193 photo-identifications of 454 individuals were recorded during 2,851 encounters throughout 1,024 days of sampling across 16 years (mean 64 days of sampling per year; SD = 32, minimum = 22, maximum = 144; Fig S1). Overall, we collected an average of 5.22 identifications per day of sampling (4.8 unique individuals d-1; 1.1% of the identified population d-1), and the mean number of encounters per individual was 11.4. The number of new identifications increased steadily from 2004 to 2016, then began to level off (Fig. S2). The rate of new identifications per encounter dropped steeply from 2004 to 2008 and has slowly leveled off since (Fig. S2), and the percent of encounters with previously identified whales increased from ∼50% in 2004-2006 to more than 90% in 2017-2019 (Table 1). These discovery metrics indicate that, although new individuals continue to be recruited, we have identified the majority of humpback whales within the local population.

**Table 1.** Capture history of humpback whales in the Kitimat Fjord System, 2004-2019.

| Year | Identifications |  |  | Recaptures |  |  |  |  |  |
| --- | --- | --- | --- | --- | --- | --- | --- | --- | --- |
|  | Total | Unique | New | Within season |  | Prior year |  | Any prior year |  |
|  |  |  |  | *n* | % | *n* | % | *n* | % |
| 2004 | 79 | 41 | 41 | 38 | 48% | - | - | - | - |
| 2005 | 168 | 38 | 18 | 130 | 77% | 20 | 53% | 20 | 53% |
| 2006 | 230 | 68 | 33 | 163 | 70% | 27 | 40% | 35 | 51% |
| 2007 | 141 | 55 | 19 | 86 | 61% | 34 | 62% | 36 | 65% |
| 2008 | 148 | 75 | 32 | 73 | 49% | 28 | 37% | 43 | 57% |
| 2009 | 297 | 96 | 33 | 201 | 68% | 46 | 48% | 63 | 66% |
| 2010 | 252 | 85 | 21 | 167 | 66% | 54 | 64% | 64 | 75% |
| 2011 | 242 | 103 | 41 | 139 | 57% | 44 | 43% | 62 | 60% |
| 2012 | 239 | 100 | 19 | 139 | 58% | 50 | 50% | 81 | 81% |
| 2013 | 452 | 151 | 52 | 301 | 67% | 68 | 45% | 99 | 66% |
| 2014 | 750 | 164 | 43 | 586 | 78% | 97 | 59% | 121 | 74% |
| 2015 | 874 | 213 | 58 | 661 | 76% | 120 | 56% | 155 | 73% |
| 2016 | 615 | 181 | 28 | 434 | 71% | 133 | 73% | 153 | 85% |
| 2017 | 113 | 77 | 4 | 36 | 32% | 58 | 75% | 73 | 95% |
| 2018 | 240 | 103 | 5 | 137 | 57% | 50 | 49% | 98 | 95% |
| 2019 | 353 | 126 | 7 | 227 | 64% | 67 | 53% | 119 | 94% |
| Mean | 325 | 105 | 28 | 219 | 62% | 56 | 50% | 76 | 68% |
| SD | 233 | 50 | 16 | 186 | 12% | 35 | 17% | 45 | 23% |

### Habitat use

#### Site fidelity

##### Interannual

The mean annual recapture rate was 50% (SD=17%, min = 37% in 2008, max = 75% in 2017; Table 1). Of the individuals we identified, 263 (58%) were seen in more than one year, 116 (26%) were seen in 5 or more years, 49 (11%) were seen in 10 or more years. Three individuals were seen in all 16 years of the study (Table 2). Across the entire study, 137 individuals (30% of identified population) were encountered on ≥ 10 occasions.

**Table 2.** Recapture statistics for identified humpback whales in our study. *Left*: The number of whales seen in at least 1, 2, … 16 years. *Right*: The number of whales seen in at least 1, 2, … 126 encounters.

| Recaptures |  |  |  |
| --- | --- | --- | --- |
| *Years* | *Individuals* | *Encounters* | *Individuals* |
| 1 | 454 | 1 | 454 |
| 2 | 263 | 2 | 315 |
| 3 | 208 | 5 | 192 |
| 4 | 146 | 10 | 137 |
| 5 | 116 | 20 | 83 |
| 6 | 98 | 30 | 49 |
| 7 | 83 | 40 | 34 |
| 8 | 68 | 50 | 24 |
| 9 | 56 | 60 | 14 |
| 10 | 49 | 70 | 11 |
| 11 | 40 | 80 | 11 |
| 12 | 34 | 90 | 9 |
| 13 | 29 | 100 | 7 |
| 14 | 20 | 110 | 3 |
| 15 | 9 | 120 | 1 |
| 16 | 3 | 126 | 1 |

##### Seasonal

On average, approximately 15% of identified whales arrived by the end of May (doy 150; Fig S2). In most years, the rate of arrival increased between late June and mid-August (doy 180 – 230). The fact that new individuals were continually identified throughout each field season, even in longer seasons greater than 200 days in length, indicates that the population is not closed within a single year.

The mean recapture rate within a season was 62% (SD=12%; Table 2), and the mean documented occupancy within the fjord system (days between first and last observation) each year was 20 days (SD=25, min=1, max=130; Table S2). Permutation tests of the Lagged Identification Rate (LIR) curve indicate that humpback whales remain in the study area for 50 days longer than expected if they were conducting random movements in and out of the study area (Fig 2; see Supplementary Appendix 1 for additional results).

**Figure 2.**
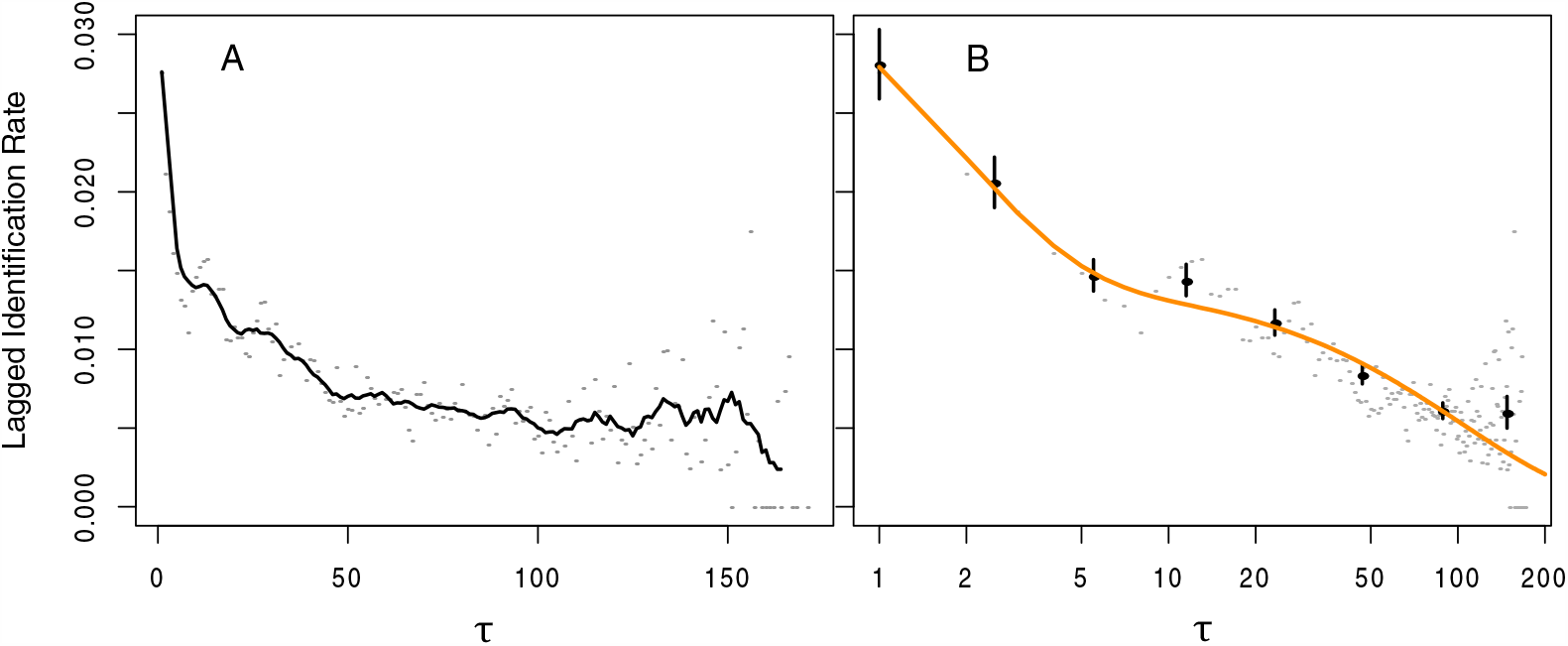
Lagged Identification Rate (LIR) of humpback whales in the Kitimat Fjord System. *Left:* Grey dots represent the LIR calculated for each time lag (*τ*, in days) tested. Black line is the running mean of LIR (window=10 days, first point forced to the *τ* = 1). Blue line and shaded area represent the median and 95% confidence interval (2.5% and 97.5% quantiles), respectively, of the permutation tests (n=100). Lags at which the running mean rises above the shaded area indicate significant patterns in residency behavior. *Right:* Best-fitting SOCPROG model of LIR (note log scale). Points with standard errors (n=100 bootstraps) are lag-pooled calculations of the LIR, computed at *τ* = 2^0―8^. Orange line represents the best-fitting model (see Table S6).

##### Bubble net feeders

The subpopulation of known bubble net feeders (n=128 individuals; 65% of whales seen 5 or more times), when compared to the remainder of the identified population, scored significantly higher (p < 0.001) in all site fidelity metrics tested: annual rate of return, seasonal occupancy, permanence, periodicity, Standardized Site Fidelity Index (SSFI), and the mean minimum stay. Bubble netters arrived earlier than other whales (p < 0.001), but there was no significant difference in the mean date of final encounter (p = 0.456), which is consistent with longer stays in the area. Based on randomization tests, the LIR of bubble net feeders was significantly greater than the remainder of the identified population for time lags of 1 − 70 days (Fig. S3).

#### Behavior

Randomization tests indicated that geographic patterns of several aspects of habitat use (Fig. S4) were significantly unlikely under the null hypothesis that whale behaviors were distributed randomly in space and time (Fig. 3). Bubble-net feeding occurred more frequently than expected in the outer channels and less frequently within inner channels; conversely, other feeding modes (including subsurface feeding inferred from surface behaviors) exhibited the opposite pattern. Consistent with the previous findings in the area [84], encounters in outer channels occurred early in the year, and encounters deep within the fjord system occurred later on.

**Figure 3.**
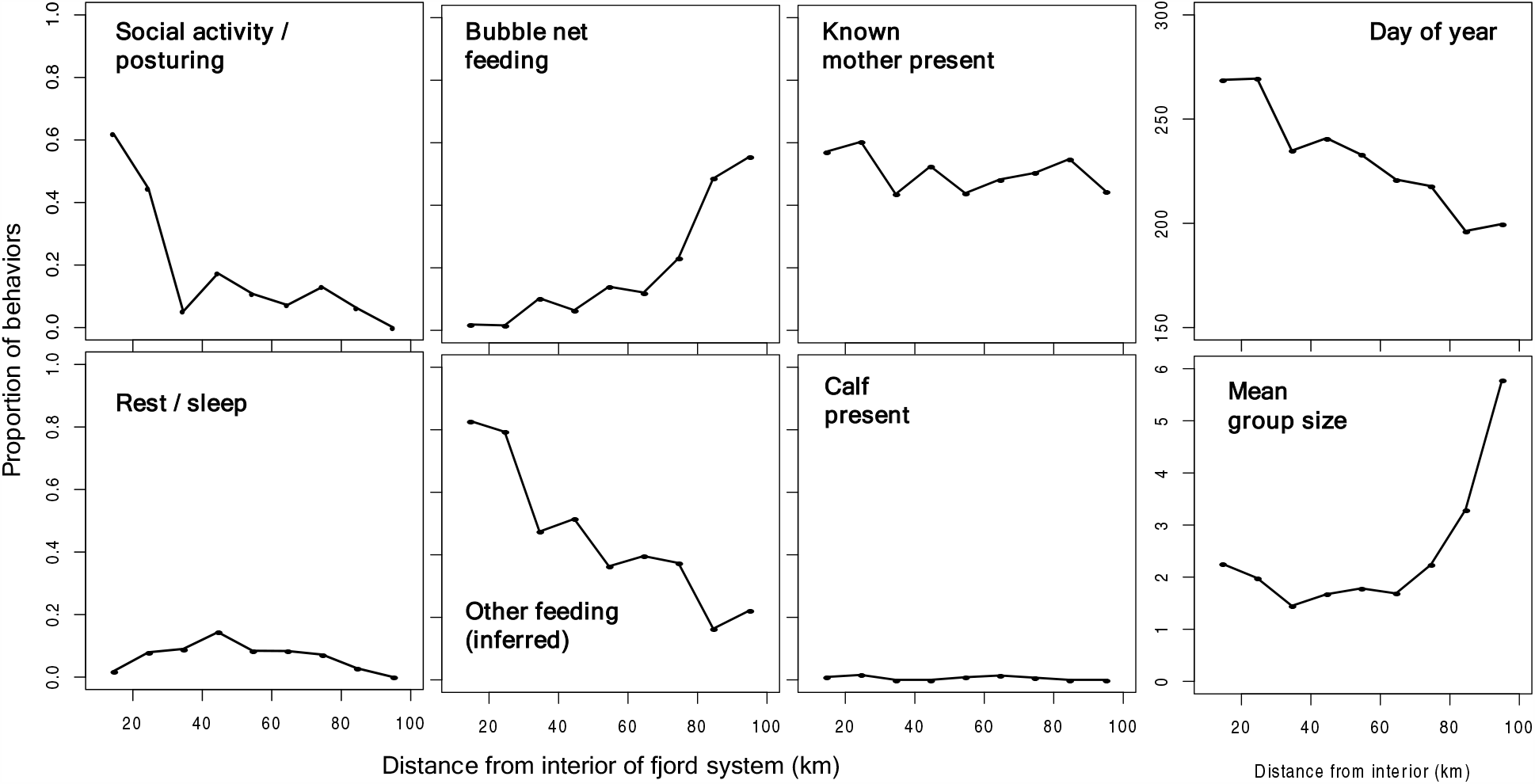
Geographic patterns in the behavior, seasonality, and habitat use of humpback whales within the Kitimat Fjord System. In each pane, the black line indicates the frequency of each variable of interest, pooled into bins of swimming distance from the interior of the fjord system (bin size = 10km, 0-100 km. Blue shaded area represents the frequency expected by random chance, determined via randomization (1,000 iterations; blue line = median; shading = 95% confidence interval, determined using 0.025 and 0.975 quantiles).

Mean group size was higher than expected in the outer channels as well as deep within the fjord system, but not in central channels. The large groups of the outer channels are attributable to bubble net feeding groups; our generalized linear mixed model (GLMM) indicated a significant difference between bubble-net group size and all other behavior categories (Table S1; Fig. S5; Fig. S6). In the deep inland channels, the groups we encountered commonly occurred within dispersed aggregations of whales (30 or more, on occasion) who, while interacting closely with only one or two individuals for intermittent periods, also exhibited more fluid associations and a higher rate of social mixing, periodically forming larger groups in which pronounced social activity and posturing, without any apparent connection to feeding, were observed. Those unusual social behaviors occurred at a significantly high rate deep within the fjord system and less than expected under random chance expectation in other portions of the study area. These unusual social behaviors occurred almost exclusively in late summer and early fall. See Supplementary Appendix 2 for additional results.

### Sociality

#### Stability of associations

Of the 454 whales we identified, 355 (78%) were observed in association with one another as dyads (Table 3). On 5 or more occasions we observed 276 of these dyads, involving 35 individuals. 22 dyads (10 whales) were observed on 10 or more occasions, and six whales were involved in 8 dyadic associations that were observed at least 35 times. Many dyads (n=908 dyads of 133 whales) were observed in multiple years (Table 3). Of these, 155 dyads (20 whales) were observed in at least 5 years, and 17 dyads (7 whales) were observed in at least 10 years. Seven dyads (6 whales) were observed in 12 years.

**Table 3.** Dyad recapture statistics for pairs of associated humpback whales in our study. *Left*: The number of dyads seen in at least 1, 2, … 16 years. *Right*: The number of dyads seen in at least 1, 2, … 126 encounters.

| Dyad recaptures |  |  |  |  |  |
| --- | --- | --- | --- | --- | --- |
| *Years* | *Dyads* | *Individuals* | *Encounters* | *Dyads* | *Individuals* |
| 1 | 4,358 | 355 | 1 | 4,358 | 355 |
| 2 | 908 | 133 | 2 | 1,146 | 151 |
| 3 | 370 | 60 | 3 | 545 | 78 |
| 4 | 217 | 30 | 5 | 276 | 35 |
| 5 | 155 | 20 | 10 | 129 | 19 |
| 6 | 119 | 17 | 15 | 68 | 13 |
| 7 | 85 | 14 | 20 | 31 | 10 |
| 8 | 63 | 14 | 25 | 22 | 7 |
| 9 | 46 | 13 | 30 | 16 | 7 |
| 10 | 17 | 7 | 35 | 8 | 6 |
| 11 | 9 | 6 | 40 | 1 | 2 |
| 12 | 7 | 6 | 71 | 1 | 2 |

Randomization tests of Lagged Association Rates indicate that, on average, dyads remained associated for two months longer than would be expected based on the null model of random association-dissociation (Fig. 4; Supplementary Appendix 3). The LAR of whales occurring in groups that contained known bubble net feeders (n=2,255 encounters) was significantly higher at greater time lags when compared to groups that contained whales not known to bubble net feed (n=943; Fig. 4). The former LAR was significantly higher than the null model up to time lags of 60 days; while the latter LAR was significant within a time lag of only 5 days. Dyadic associations in which at least one individual was known to bubble net feed were observed in more encounters and in more years than dyads in which neither whale was known to bubble net feed (Kruskal-Wallis rank sum test, χ^2^ = 192, df=1, p < 0.001).

**Figure 4.**
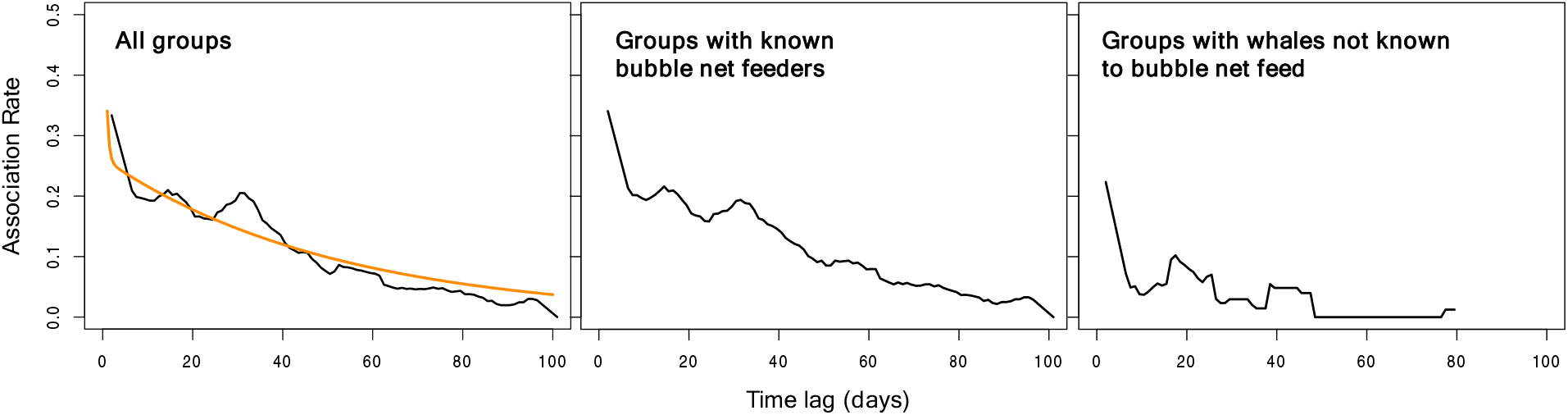
Lagged Association Rate (LAR) of humpback whales in the Kitimat Fjord System, parsed into three subsets of our data: all groups encountered (*left*), groups containing known bubble net feeders (*center*), and groups containing whales not known to bubble net feed (*right*). Grey dots represent the Association Rate calculated for each time lag (in days) tested. Black line is the running mean of LIR (window=10 days, first point forced to the *τ* = 1, last point forced to the final *τ*). Blue line and shaded area represent the median and 95% confidence interval (2.5% and 97.5% quantiles), respectively, of the randomized null model (n=100). Lags at which the running mean rises above the shaded area indicate significantly stable dyadic associations. The orange line on the left plot (‘All groups’) represents the best-fitting SOCPROG model of LAR (see Table S7).

#### Association network

The association network was moderately connected, realizing 14% of the total number of possible dyadic associations within the identified population (Table S3). The population exhibited strong social differentiation (*S* = 9.9, *r* = 0.92), providing sufficient power to test the hypothesis that KFS humpback whales had no preferred or avoided relationships (*S*^*2*^ *x H* = 9.9^2^ x 11.4 >> 5; [94]). The SRI of most association pairs (77%) fell between 0.01 and 0.05 (median = 0.021; mean = 0.028; SD = 0.024; Table S3). The highest SRI value between any dyad was 0.438. Based on data stream permutations, strong associations were more common than would be expected from random association-dissociation dynamics (p < 0.001). 11% of non-zero associations were stronger than expected from chance at α=0.05.

#### Social preferences

We used Generalized Affiliation Indices (GAIs) to control for the influence of non-social factors (i.e., co-occurrence in space and time and individual gregariousness, Table S4) in our understanding of the humpback whale social network (Table S3). The majority of dyadic GAIs (87%) were negative, which may be indicative of social selectivity and/or avoidance.

According to data stream permutations, the prevalence and strength of social preference fell short of significance (p=0.088) in tests of the entire population. However, the significance of social preferences differed for subsets of the population. Within the subnetwork of known bubble net feeders (5,420 dyads of 108 whales), social preferences were more common than expected (p=0.006) and stronger than expected, with borderline significance (p=0.057). The strongest GAI value among bubble net feeders was 9.05. In contrast, for the remaining subnetwork of whales not observed to bubble net feed (1,052 dyads of 50 whales), social preferences were less common (p=0.800) and weaker (p=0.924) than expected, the strongest GAI was lower (3.542), and a larger percentage of GAIs were negative (94%; compare to 80% within the subnetwork of bubble net feeders). Social preferences were similarly weak for dyadic associations between known bubble net feeders and whales not known to bubble net feed (4,867 dyads of 158 whales; Table S3).

#### Social network structure & stability

We found support for subdivision of the observed social association network into communities of individuals with stronger in-group relationships. Seven communities were identified within the network (no multi-level or nested clusters; modularity = 0.3, where modularity is a measure of the strength of network structure; Fig 5). Randomization tests indicated that the network modularity was significantly greater than would be expected based upon random network clustering (p = 0.000; mean modularity of randomized network = 0.229, sd = 0.003, min = 0.218, max = 0.241), and the observed number of communities was significantly lower than expected (p = 0.002, mean of randomization set = 10.278, sd = 0.964, min = 7, max = 13).

**Figure 5.**
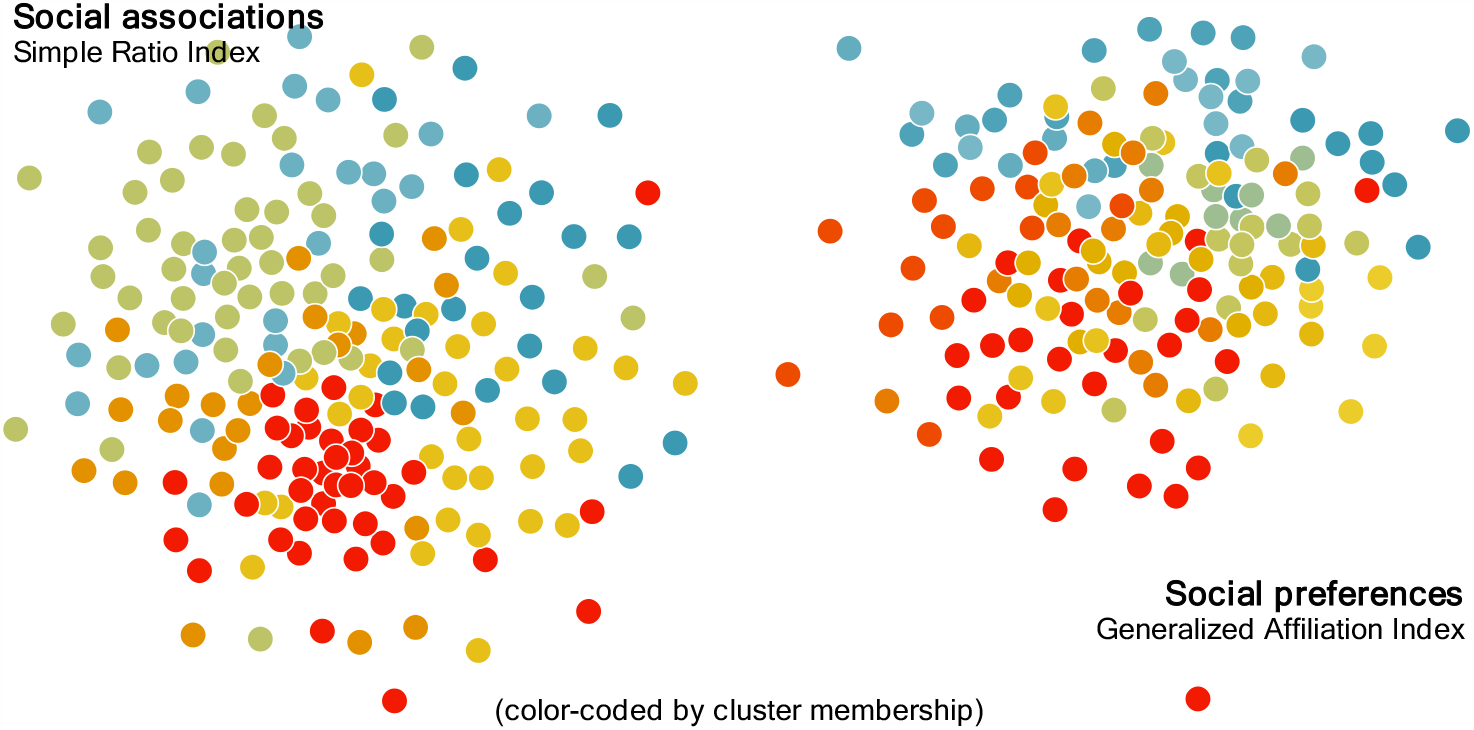
Social networks, color-coded by community membership as identified by the Louvain clustering algorithm in *igraph*, of humpback whales (≥ 5 encounters) in the Kitimat Fjord System. The left network is based upon social associations (weighted by the Simple Ratio Index) and contains 7 identified communities. The right network includes the network of social preferences (weighted by the Generalized Affiliation Index), with 13 identified communities.

One of these seven communities consisted of a single whale who was encountered on five occasions in a single year, but never with another whale. Within the remaining six communities, social differentiation ranged from moderate to strong (range in S = 0.55 – 1.77). Social differentiation appeared to be related to the proportion of dyads in which both whales were known to bubble net feed (linear regression, n=6, p=0.02, r^2^=0.77). The most differentiated community contained a high proportion (0.85) of bubble net feeder dyads, while the least differentiated community contained the lowest proportion (0.24).

The structure of the association network shifted roughly halfway through the 16-year study (Fig. S7). In the early years (2004 – 2011), the humpback population was dispersed into a higher number of smaller communities than would be expected by chance (Fig. 6A-B), with the exception of a core network of regular bubble net feeders (Fig. S7). In contrast, during the more recent years of 2012-2019, network connectivity increased such that the number of discrete communities declined and the five largest contained a larger proportion of the population than expected (Fig. 6A-B). During this 2010-2012 shift, community modularity broke down and was similar to what would be expected from random social mixing (Fig. 6C).

**Figure 6.**
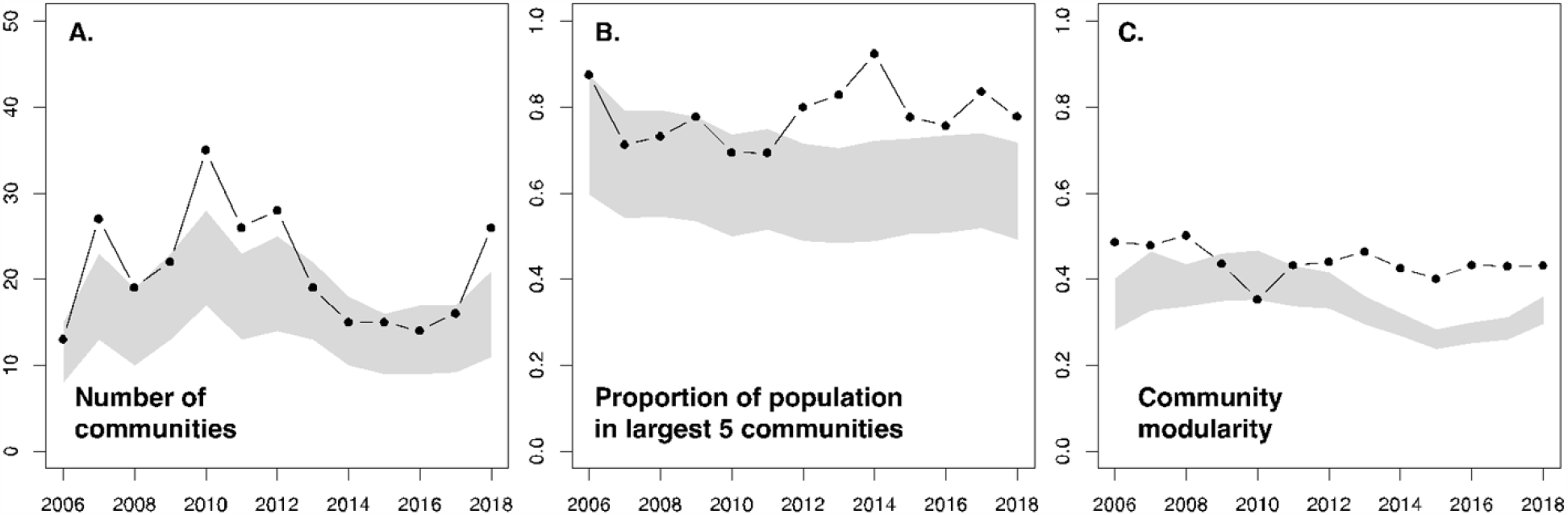
Annual time series of association network structure metrics, based on a four-year running window (2004-2007, 2005-2008, …, 2016-2019; n=12) for the population of humpback whales seen on at least five occasions throughout the 16-year study. The observed structure metrics (black line and dots) are compared to a null model based upon 1,000 randomizations each of the original sighting records within each four-year interval (grey shaded area is 95% confidence interval of these randomizations, based on 0.025 and 0.975 quantiles). Significantly non-random structural patterns are indicated by the departure of the black line from the grey shaded area. **A**. The number of communities within the network, as identified by the Louvain algorithm in igraph. **B**. The proportion of the population contained within the five largest communities identified by the clustering algorithm. **C**. The modularity of the community structure.

### Relationship between social structure and habitat use

#### Dyadic associations

Engaging together in certain behavioral contexts was correlated strongly with engaging together in others (Table 4). Dyads who were seen sub-surface feeding together were more likely to travel together (p=0.000) compared to those that were not, and, conversely, dyads found traveling and exhibiting robust behavior were more likely to be found sub-surface feeding (p=0.004 and p=0.002, respectively). Bubble net feeding together, however, did not, on average, translate to associations in other behavioral contexts. Bubble net feeding dyads were significantly unlikely to be found engaged in sub-surface feeding (p=0.000) or resting (p=0.004).

**Table 4.** Social transference of behavior. The probability that humpback whales dyads engaged in a primary behavior would ever be observed on a separate occasion to be engaged in a secondary behavior, as determined by randomization tests (n=1,000; see main text) applied to the subset of dyads seen together on at least three occasions. Significant findings, based on a two-tailed significance test (p < 0.025 or p > 0.975) are in boldface. Column *n* provides the sample size of the dyad sets used in the test; the first number is the size of the dyad set known to practice the primary behavior together, and the second number is the size of the set of dyads who were not.

| Primary behavior | $n$ | Secondary behavior | | | | |
| --- | --- | --- | --- | --- | --- | --- |
|  |  | Bubble net | Feed (other) | Travel | Robust | Rest |
| Bubble net | 336, 35 | - | **0.000** | 0.872 | 0.075 | **0.004** |
| Feed (other) | 153, 218 | **0.001** | - | **1.000** | 0.940 | 0.901 |
| Travel | 210, 161 | 0.942 | **0.996** | - | 0.517 | 0.451 |
| Robust | 29, 342 | **0.001** | **0.998** | 0.523 | - | 0.630 |
| Rest | 39, 332 | **0.000** | 0.961 | **0.998** | 0.783 | - |

#### Association networks

Figure S8 shows the network of associations in the context of individual-level patterns in habitat use and behavior. These visualizations highlight potentially related patterns of social assortment within the network. For example, bubble net feeding whales are assorted predominantly into the lower half of this representation of the network. Those same individuals, when compared to the remaining population, appear to exhibit greater site fidelity, arrive earlier in the year, occur predominantly in the outer channels of the fjord system, and spend less time conducting explicitly social behaviors such as ‘posturing’. Other traits, such as being a known mother (Fig. S9), did not appear to be assorted within the network. These patterns are verified statistically below.

Of the 63 behavioral k-means clustering schemes we compared to the social clustering, 11 (17%) had statistically significant (p < 0.05) congruency indices based upon comparison to a null distribution of randomized social associations. All significant clusterings yielded a p-value of 0.014 or less. The most congruent clustering (p=0.000; adj. rand = 0.027) was based upon the bubble net feeding rate and the standard deviation of distance from the fjord interior. The significance of this variable set was confirmed by reverse randomization, in which the observed congruency of the social-behavioral clusterings was greater than expected based on a null model of random behavior (p=0.004). All six behavioral variables tested were present in at least one clustering in the significance set. The standard deviation of fjord position was most often included in significant clusterings (73%), followed by resting rate (64%), then bubble net feeding rate (55%).

Of the 15 site-fidelity k-means clustering schemes we compared to the social clustering, 8 (53%) were significantly congruent. All significant clusterings yielded a p-value of p=0.015 or less. The most congruent clustering (p=0.000; adj. rand = 0.036) was based upon number of years present, minimum stay, and the Standardized Site Fidelity Index (SSFI) (reverse randomization confirmed significance; p=0.001). All four variables tested were present in at least one clustering in the significance set, and number of years present was included in all significant clusterings.

Randomization tests indicated that behavior and site fidelity clusterings based on k=7 communities were also significantly congruent (p < 0.001). When behavior and site fidelity variable sets were combined (n=1,023 combinations), 50% of sets yielded significantly congruent (p ≤ 0.05) community assignments, and 92 (9%) were congruent at p < 0.0001. Within these variable sets, the five most frequently included variables were years seen (99% of variable sets), SSFI (70%), standard deviation of fjord position (68%), bubble net feeding rate (59%), and social rate (56%).

#### Social preference networks

When mapping individual-level traits upon the network of social preferences (Fig. 7), the assortment patterns described above remained evident. Based upon assortativity coefficient (AC) significance testing (Table S5) for many of the traits tested, trait similarity between individuals appeared to reduce avoidance and facilitate social assortment. This is indicated by lower ACs than expected within the negative GAI network for the following traits: bubble net feeding rate (p=0.000), feeding rate (p=0.001), social rate (p=0.001), mean fjord position (p=0.000), SD fjord position (p=0.004), and average arrival date (p=0.0001). Of these, feeding rate also registered as a significant assortment factor among preferred relationships (p=0.007), and bubble net feeding rate and mean fjord position fell just short of significance (p=0.075 and p=0.040, respectively).

**Figure 7.**
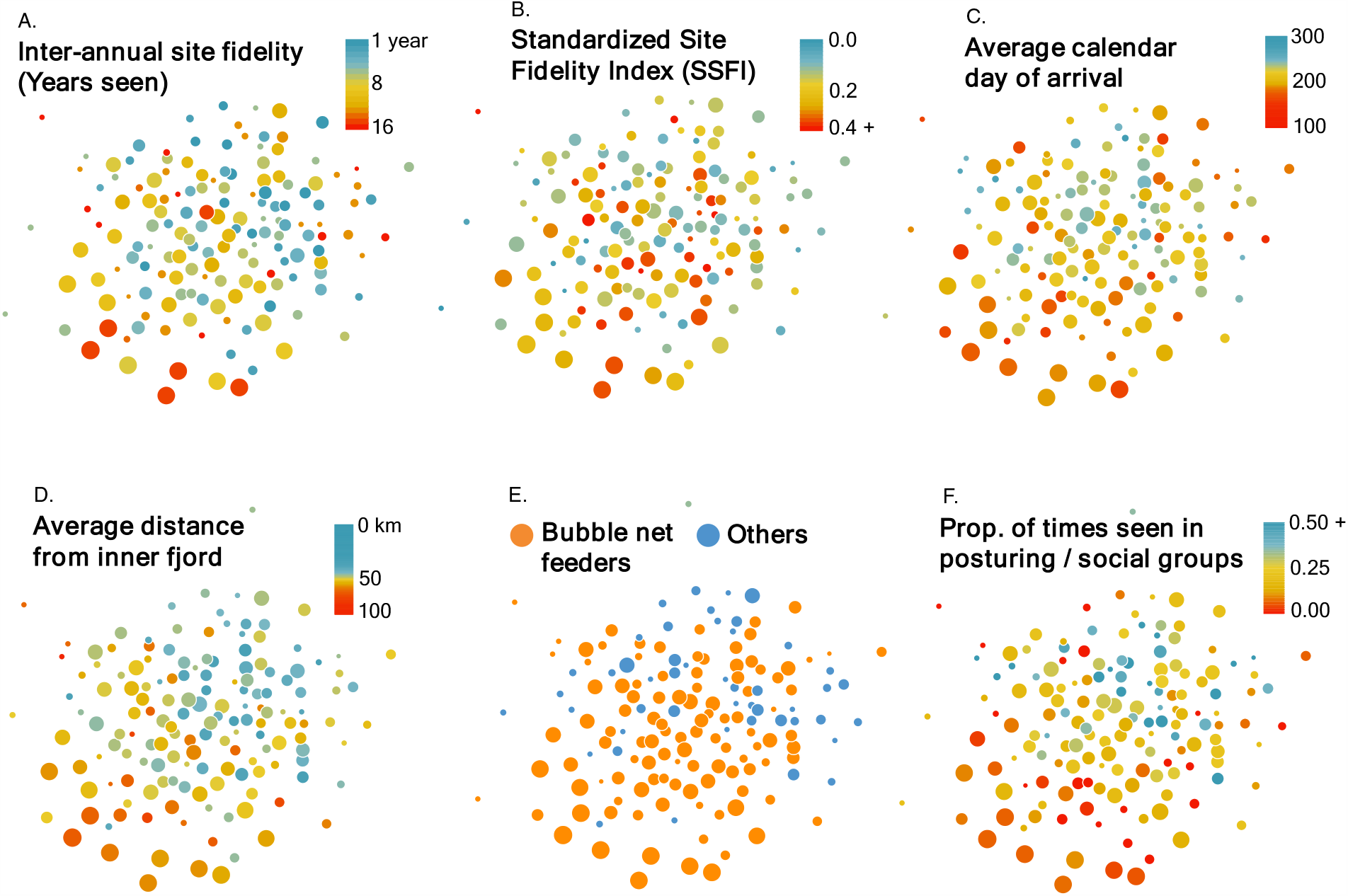
Network of social preferences of humpback whales (≥ 5 encounters; n_edges_ = 1,527; n_whales_ = 136) based upon the Generalized Affiliation Index and color-coded by various aspects of habitat use and behavior. In all networks the placement of individuals remains the same, demonstrating related patterns in the distribution of individual traits. Vertex size reflects the number of observations of each individual. Thicker, darker edges represent statistically significant associations based upon data stream permutations (1,000 iterations). Networks built with igraph in R using the Fruchterman-Reingold layout.

Interestingly, two site fidelity traits (proportion of years seen and average minimum stay) yielded significantly large ACs within the negative GAI network (p=1.00 and 0.994, respectively), and the AC for the categorical variable of ‘known bubble net feeder’ (yes or no) was nearly significant (0.900; Table S5). These traits appear to be important factors for avoidance, in that whales who are dissimilar in these respects tend to avoid social interaction, even when controlling for structural variables such as geographic and temporal overlap.

Network position ∼ trait correlations also indicated that strategies of habitat use mediated the network of social affiliations in ways consistent with our other findings (Table 5; see Table S8 for corresponding analysis using association indices instead of affiliation indices). Whales known to bubble net feed exhibited significantly higher degree-, betweenness-, and closeness-centrality and had more contacts than expected (p < 0.001). Similar network position patterns were seen in whales with high annual rates of return, and within-season residency (SSFI) was also significantly correlated to high centrality (p < 0.01). Conversely, the position ∼ trait correlation was significantly weaker than expected for whales that commonly practiced other modes of feeding, and we found no correlation between network positions and social rate, resting rate, or status as a known mother. Whales who were encountered primarily in offshore channels exhibited high degree- and closeness-centrality (p ≤ 0.002), while whales whose fjord position varied greatly exhibited high betweenness centrality (p=0.004). Whales that remained in the area for longer stays exhibited higher degree centrality (p=0.003) and a higher number of contacts (p=0.001).

**Table 5.** P-values indicating the relationships among affiliation network position metrics (based on positive GAIs) and strategies of behavior, habitat use, and site fidelity in the population of humpback whales in the Kitimat Fjord System. P-values represent the proportion of randomizations for which the slope coefficient of the regression (Position metric ∼ Trait) that is greater than the real observed value. Based on a two-tailed test with α=0.05, P-values below 0.025 (bold) indicate that the relationship between the position metric and the trait is significantly stronger than expected. P-values (also bold) that are greater than 0.975 indicate that the relationship is significantly weaker than expected.

| Category | Trait | Individual network position metric |  |  |  |
| --- | --- | --- | --- | --- | --- |
|  |  | Degree centrality | Betweenness centrality | Closeness centrality | Contacts |
| Behavior /<br>Biology | Known bubble netter | **0.000** | **0.000** | **0.000** | **0.000** |
|  | Bubble net rate | **0.000** | **0.003** | **0.000** | **0.000** |
|  | Feeding rate | **1.000** | 0.901 | **1.000** | **1.000** |
|  | Social rate | 0.964 | 0.117 | 0.492 | 0.808 |
|  | Resting rate | 0.990 | 0.714 | 0.991 | 0.892 |
|  | Known mother | 0.214 | 0.441 | 0.771 | 0.030 |
| Habitat use | Mean fjord position | **0.000** | 0.392 | **0.002** | 0.027 |
|  | SD fjord position | 0.189 | **0.004** | 0.131 | 0.093 |
| Site fidelity | Years seen | **0.000** | **0.001** | **0.000** | **0.000** |
|  | SSFI | **0.009** | 0.111 | **0.005** | 0.033 |
|  | Average arrival date | 0.984 | 0.138 | 0.594 | 0.956 |
|  | Minimum stay | **0.003** | 0.409 | 0.239 | **0.001** |

## DISCUSSION

Through long-term study of humpback whales within an important feeding habitat, we have observed complex interactions between patterns of habitat use and their social lives. By identifying modes of habitat- and ecological-partitioning and finding that such traits correlate strongly to the social structure of the population, we have presented evidence of social niche partitioning on the feeding grounds of a baleen whale. As with similar findings in populations of odontocetes (e.g., [31,38,45,57,59,60]) and baleen whales in breeding areas (e.g., [28,115]), the complexity of habitat use and population structure exhibited here has implications for our understanding of humpback whale ecology and conservation.

### Habitat use

Strong site fidelity is a commonly recognized feature of humpback whale populations, particularly with respect to feeding grounds [116], and has direct bearing on how whale habitat is prioritized and managed for conservation [117]. Individuals within discrete feeding aggregations return to the same regions year after year according to maternal lineage and imprinting [61,63,118,119]. Even in discrete areas *within* the range of a feeding aggregation, strong site fidelity to particular subregions has been observed [120].

The humpback whales of the Kitimat Fjord System (KFS) belong to the Northern British Columbia – Southeast Alaska feeding aggregation (NBC-SEAK, [121]), and the vast majority of them (85% - 94%, [122,123]) migrate to breeding grounds in Hawaii and thus belong to the Hawaiian Distinct Population Segment [124]. The return rates we observed (mean 50%, range 37% - 75%) are similar to those documented elsewhere within this feeding aggregation, e.g., 47.2% in the southeast coastal Alaska [125] and 63% in Glacier Bay, Alaska [126]. Further to the northwest in Alaska, mean return rates have ranged from 23% (western Aleutians; [127]) to 34% (Kodiak; [128]) to 37% (Shumagin Islands, AK; [128]). On feeding grounds in other ocean basins, documented return rates include means of 41% (Gulf of Corcovado, Chile; [129]) and 73% (Gulf of Maine, USA; [130]). On breeding grounds, interannual site fidelity appears to be substantially lower, both in the north Pacific [116,131] and elsewhere in the world [132,133]. Unfortunately, the annual return rates and occupancy durations documented within site fidelity studies can be biased by several factors, ranging from survey effort to population size to sex ratios to fluking behavior [128,130,134]. Hence the rates observed across studies and regions can only be compared loosely. Nevertheless, our findings confirm the importance of seasonal residency to the way humpback whales use habitats within the NBC-SEAK feeding aggregation area.

The seasonal occupancy we observed in the KFS (mean = 20 d) was lower than elsewhere within the NBC-SEAK feeding aggregation, such as Glacier Bay (mean residency duration = 67 d; [126]), and falls in between observations from other ocean basins, e.g., 15 d in the Gulf of Corcovado, Chile [129] and 88 d in the Gulf of Maine [130]. Occupancy at breeding areas tend to be much shorter; documented averages include 5 - 15 d off Brazil [132,133], 6 – 9 d in the Caribbean [76,89], and 13 d in Ecuador [135]. However, longer breeding ground occupancies have been observed (e.g., 34 d in Hawaii; [136]; 40 d off Camiguin Island; [137]). The intermediate occupancy we observed on the foraging ground we studied, if not an artefact of methodologies or behavior, may reflect the fact that the KFS is located more or less at the geographic center of this feeding aggregation’s range, and therefore may be more likely to be visited while individuals transit between feeding grounds to the north and south. If this is the case, then the fjord system’s location may be a contributing factor to the complexity of habitat use strategies we have observed within it.

The humpback whales of the KFS exhibit a wide array of habitat use strategies, some of which were associated with strong social interactions. At one end of this spectrum are individuals who perform brief, one-off visits to this habitat with no clear preferences in spatial use, prey, or social affiliation. At the other end are seasonally resident individuals who return on an annual basis, or nearly so, and who have strong tendencies in their choice of habitat features, prey type, feeding mode, and social group. Between these two extremes, the varieties of strategies vary in four salient ways: primary mode of feeding, habitat selection within the fjord system, the seasonal timing of their residency, and the variety and stability of their social bonds. We found that these strategies were all correlated; for example, bubble net feeders tend to use the outer channels of the fjord system, arrive early in the year, stay in the area for longer periods, and exhibit stronger social preferences and greater social selectivity. Other individuals, in contrast, did not engage in bubble net feeding at all, appeared to practice deep-feeding behavior (likely targeting krill; [138]), occurred more commonly deeper into the fjord system, arrived later in the year, were more socially fluid, but engaged in more purely ‘social’ behavior in the sense that their interactions were often not tied in any apparent way to feeding. The fact that these social niches are manifest along a continuum differs from examples of social niche partitioning observed in odontocetes such as sperm whales (e.g., [45]) and killer whales (e.g., [57]), and may reflect the fact that this feeding habitat was only recently reoccupied by humpback whale populations that are still in the process of recovering, in terms of both abundance and ecological role, from commercial whaling [81,108]. Furthermore, it is unknown whether or not the social and ecological roles manifest within this population will recover in step with its abundance, if at all, nor is it known whether or not the patterns emerging in the KFS even reflect those that existed before and during commercial whaling. Future years of study within this habitat will allow us to interpret the stability and discreteness of the social niches we have documented here.

The social niche partitioning we report here offers further insight into previous studies of habitat use within this same humpback whale population and study area. Keen *et al*. [84] observed that humpback whales occupied the KFS in a ‘wave’ pattern, in which whale distribution is concentrated in offshore channels in the early summer, then propagates to deep inland channels in the autumn. Active acoustic surveys were used to relate prey distribution to this ‘whale wave’, but that correlation was found to be very weak, particularly in the autumn months, and the authors were ultimately unable to determine the drivers of this strange habitat use pattern. That study was based on a subset of the data we have presented here, but it relied only upon the locations of whale encounters, and did not include photo-identifications. Here, individual sighting histories and social records have demonstrated that the ‘whale wave’ within this fjord system is, in fact, the result of social niche partitioning. While the distribution of whales does shift inland throughout the summer, we can now say that the composition of individuals is also turning over, and that these individuals exhibit specific habitat preferences within the fjord system. In general, early arrivals exhibit a preference for the fjord’s outer channels, late arrivals prefer inner channels, and intermediate arrivals either prefer the central channels or have weaker preferences. With this insight in hand, now the most glaring gap in our understanding of these humpback whales is the genetic structure of their population, and whether or not it corresponds to the patterns of social structure and habitat use we have presented here.

The importance of bubble net feeding as an organizing factor in this population was reiterated throughout our findings. Compared to the remainder of the population, individuals who engaged in bubble net feeding exhibited higher site fidelity to the fjord system, stayed in the area for longer durations, were involved in longer-lasting, stronger social relationships, exhibited strong social preferences, were more socially connected and exhibited greater centrality within the social network, and were members of social communities that exhibited higher social differentiation. Finally, bubble net feeding rate was an important explanatory variable in the overlap of social clusters and modes of habitat use. Clearly, social niche partitioning within this population is mediated strongly by bubble net feeding behavior.

Humpback whales frequently forage in groups, targeting a variety of prey using a diverse repertoire of feeding strategies [34,65,139–141], but bubble net feeding is remarkable in its complexity and spectacle [142]. Now known to require sophisticated coordination involving the division of labor and role specialization ([35]; but see observations of solitary bubble netting in our Supplementary Material), bubble net feeding has been observed in only a few subpopulations targeting only certain prey: herring (*Clupea harengus*) in southeast Alaska [34], sand lance (*Ammodytes spp*.) in the southern Gulf of Maine [139,142], and Antarctic krill (*Euphausia superba*) in the Southern Ocean [35].

In the KFS, bubble net feeding is similar in form to that described in southeast Alaska, in which groups of whales dive below a school of herring, release spiraling curtains of bubbles that corral the fish, ascend steadily until the fish are trapped at the surface, then lunge vertically through the school [34,140,143]. During a bubble net feeding event, one individual (or in rare cases more than one; authors, unpubl. data) also produces a distinct tonal bellow that has come to be known as a ‘feeding call’ [34]. Although the KFS is the only location in Pacific Canada where this form of bubble net feeding has been scientifically documented [80,84,85,138], we have observed it elsewhere in Canadian waters: further north near Prince Rupert (N 54.3, unpubl. data), and as far south as Cape Caution (N 51.1, unpubl. data). As a behavior that allows humpback whales to access a prey type that individuals cannot capture effectively on their own, bubble net feeding is likely of critical importance to the species’ ecological niche, carrying capacity, and resilience to environmental perturbations within Pacific Canada [80]. In order to better understand the importance of bubble net feeding to humpback whales along this coast, on both local and regional scales, we emphasize the need for directed research regarding the underlying social dynamics of the behavior in Canadian waters, as well as its habitat requirements, particularly with respect to its sensitivity, if any, to disturbances from anthropogenic noise, fishery interactions, and nearby vessel activity.

### Sociality

Social behavior in baleen whales is typically associated with their breeding grounds [78], where most social studies have been carried out (e.g., [25,78,89,130]). As with other mysticetes, humpback whales have been characterized as largely solitary, occasionally engaging in short-lived and fluid group interactions [70,71]. On feeding grounds, humpback whales are typically found alone or in groups of two or three individuals that are spatially segregated; large groups are a rare exception, and when they occur they are typically associated with cooperative feeding [71,72,74,90,140,144,145]. Larger groups are also known to occur on breeding grounds in association with aggressive male competition [88,146,147].

Regardless of group size, group composition has been characterized as unstable and fluid, rarely lasting more than a few days [70,71,73], particularly on breeding grounds where the only stable social bonds occur between mother-calf pairs [75–77,144,148]. These short-lived bonds are not necessarily unimportant [73], and they have been found to reflect matrilineality and kinship [68,141], but they do influence our understanding of the species’ natural history and its needs within a management context.

The first evidence of long-term social bonds actually came from the first decade of humpback social studies, specifically from observations of a bubble net feeding group targeting herring in southeast Alaska [74,88,144,149–151]. Multi-year pair bonds were also observed in the early decades of social study [72,90]. As stated previously, these findings were interpreted as rare exceptions to the rule. Recent studies from feeding grounds, however, have presented compelling evidence that the social structure of humpback whales is more complex than previously thought [77,78]. In the Gulf of St. Lawrence, long-term pair bonds (up to 6 years) have been documented, with sex and reproductive status identified as factors in their temporal stability [77]. Similar findings have come from the Gulf of Maine, with the additional insight that this humpback whale society actually contains internal structure (i.e., divisible local communities; [78]).

Our findings, which include community structuring and, to our knowledge, the most enduring multi-year pair bonds on record for this species (12 years, 75% of study years), lend further evidence to this emerging picture that on feeding grounds, humpback whale society is more complex than previously thought. Examples of this now span both the Atlantic and the Pacific basins. However, our findings in the Pacific differ from those in the Atlantic in one important respect. Unlike in the Atlantic (see [77,78]), the strongest and most enduring associations we observed were not between reproductively active females, but between bubble net feeders. To fully understand the biological significance of this difference, and its implications for conservation, more social research is needed elsewhere in the feeding areas of the northeast Pacific, as well as in other ocean basins. With ecological interactions rather than demographic dynamics as a primary determinant of social structure, this Pacific population may be vulnerable to different environmental perturbations than those that have been studied in the Atlantic.

The shift in the humpback social network structure that we observed throughout the study is noteworthy for its coincidence with environmental changes and demographic shifts within the same time frame. Halfway through the study (2010 – 2012), we saw the network structure change from a fragmented community with a large number of relatively small social clusters to a more strongly interconnected community with a few disproportionately large social clusters (Fig. 6, Fig. S7). This corresponds to the same years in which the trend in this population’s calving rate shifted from a positive rate to a negative [108,152]. In the years that followed, humpback whales in the northeast Pacific Ocean experienced a series of large-scale environmental perturbations, including a debris field from the 2011 Tohoku tsunami with unprecedented levels of anthropogenic materials, a 2014 flip in the Pacific Decadal Oscillation from a strong negative phase to a strong positive phase, a strong El Niño Southern Oscillation event in 2015, perennial sea surface temperature rise, and an anomalously persistent ridge of high pressure that was established in the northeast Pacific from December 2013 to November 2015 [152–155]. These events have been implicated in food supply shortages and mass mortalities in other marine tetrapods [156,157], and may have impacted humpback whale carrying capacity within northeast Pacific feeding grounds [152].

It is possible that these environmental developments could be related to the shift we observed in social network structure within the KFS. Perhaps the increased social connectivity was in response to a lapse in resource dependability, which can force individuals to explore other habitats and strategies of habitat use, hence interacting with a larger number of individuals in the process. However, we lack the prey data necessary to parse the exact mechanics behind the changes we observed. Resource availability and stability are known to influence social structure in invertebrates [158], primates [159,160] and killer whales [161,162], but the nature of that influence depends on specific natural histories. Alternative hypotheses, unrelated to resource access, are also viable. For example, perhaps the years of 2010 – 2012 marked a shift in social structure driven simply by the density of humpback whales within the fjord system, which had been increasing since we first began studying them in 2004 (Fig. S1). Connectivity remained high later in the study even as encounter rates dropped, but it is conceivable that the social connectivity, having been established, was preserved even as population density returned to low levels. Clearly, further work is needed in this area of research. The relationship between baleen whale social structure and resource access, if better understood, could increase the conservation value of social observations in cetacean studies.

### Implications for conservation

Socially-mediated modes of habitat use, particularly those involving learning and culture, should be of intrinsic interest to conservation biologists working towards the preservation of biodiversity for its own sake [38,79,163]. But there are also practical reasons for their importance. First, such forms of habitat use involve survival skills and strategies that are “fundamental to the daily lives of these animals” [38]. Without those skills, or the social scaffolding that sustains them, threatened populations would experience even greater risk of decline. Second, socially discrete subpopulations in the same habitat may face different threats, or may respond to the same threat in different ways [38,164–166]. Third, social cohesion and cultural knowledge add to group resilience and may reduce the impacts of anthropogenic threats [38]. For these reasons, social habitat partitioning has been an important consideration in habitat protection for odontocetes, such as killer whales [57] and false killer whales (*Pseudorca crassidens*; [167,168]).

Furthermore, socially-mediated habitat use can improve resource utilization for a threatened population. Ecological studies have established that niche partitioning, in addition to enhancing biodiversity and ecosystem function, can increase a population’s access to resources [4]. When coupled with social behaviors that unlock otherwise inaccessible resources, as bubble net feeding upon schooling fish does for humpback whales who cannot capture such prey sufficiently without such techniques [35], niche partitioning becomes an even more resource-efficient habitat use strategy. In this way, social niche partitioning can govern not only *how* a population uses a habitat, but also *how much* of its resources can be used – thus increasing the value of specific habitats even more.

Socially-mediated resource-use efficiency becomes increasingly important as the extent and quality of a species’ habitat decline. Viable whale habitat is threatened worldwide by the expansion of coastal urban areas, which are associated with high rates of vessel traffic, increased ocean noise, and concentrated contaminants [169]. Urbanization is occurring within the broader context of increasingly severe biosphere perturbations such as ocean warming, acidification, deoxygenation, plasticization, and other forms of pollution [170]. These changes are steadily impinging upon areas that were once relatively pristine and remote, such as the Kitimat Fjord System [171], and areas that have been officially designated as critical habitat and yet remain unprotected [172–178].

As the availability and quality of habitat decline, it becomes increasingly urgent to effectively protect what remains. To do so, management plans should include measures that preserve each specific habitat use strategy practiced within the population, and should support ongoing research that is focused upon those site-specific strategies as well as the social dynamics that bring them about. We also emphasize that (1) the exact form of habitat protection should preserve the social processes associated with the full variety of habitat use strategies, and (2) impact assessments of potential human activities within critical habitats should consider how the social structure of the population might be affected [79].

These management considerations are directly relevant to the KFS, a proposed critical humpback habitat area facing profound alterations in the next several years. At the head of the KFS is the port of Kitimat, the epicenter of western Canadian Liquified Natural Gas (LNG) project proposals [179,180]. The new LNG Canada development, already under construction in Kitimat, is slated to add up to 700 additional annual transits of deep-sea ships (up to 122,000 DWT; [181]). If all the Kitimat shipping projects currently engaged in the environmental review process are completed according to their original timelines, large vessel traffic through this whale habitat will increase by 13.5-fold, from 112 transits in 2013 to 1,516 by 2021 [181]. This increase in ship traffic will precipitate a number of potential threats to whales, including increased pollution and risk of vessel strikes [182]. Of these, increased ocean noise is the perturbation most likely to affect the social cohesion of humpback whales in this habitat [183,184]. Of particular concern is the fact that the route of the proposed shipping lane would effectively bisect the fjord system, establishing a significant perturbation that divides the outer and inner zones of humpback whale habitat.

In order to understand the potential impact of increased noise on the social dynamics of this population, and its implications for habitat use and population viability, we suggest that the Government of Canada, Transport Canada, and BC First Nations who contribute to the decision-making and policy development in support of recovery planning for cetacean species at risk collaborate with biologists on the following research priorities: (1) Do shipping lanes affect the movements and social connectivity of baleen whales within a specific habitat? (2) Does increased noise impact the stability and frequency of pair bond interactions among baleen whale individuals? (3) How will proposed increases in shipping levels impact the hearing range and acoustic connectivity of baleen whale herds [185]? (4) Does vessel noise disrupt or alter bubble net vocalization behavior on any timescale (i.e., for a single feeding event or across years)? (5) Does vessel noise disrupt or alter the behavior, habitat use, or population viability of euphausiids or schooling fish in the northeast Pacific? (6) What ecosystem services are associated with whale habitat use in its current form and complexity, and what is the value, monetary or otherwise, of those services [186]? And (7) do fine-scale site fidelity and social associations within northeast Pacific feeding aggregations reflect kinship, as has been found elsewhere [120]? If so, the disruption or loss of humpback habitat on the local scale may be a more serious threat than previously thought. Until these questions are answered, managers should adhere to the precautionary principle [187,188] in the changes they allow within baleen whale critical habitat areas, including the Kitimat Fjord System.

## Conclusion

Along the north coast of British Columbia, we have found that the social bonds and community structure of humpback whales mediate site-specific strategies of habitat use. The converse has also been true, particularly in the case of innately social behaviors such as bubble net feeding. This reinforcing dynamic between sociality, habitat use, and place, which we have referred to here as social niche partitioning, highlights the inextricability of these whales’ ecological and social lives [79], contributes to an emerging picture of humpback natural history that is more richly social than previously thought [78,127], and emphasizes the need to study baleen whale sociality for a more complete understanding of their habitat requirements and their vulnerabilities to potential threats. In a population such as this, social disruption by human activities may be an unanticipated pathway for compromising the protection of important habitat. Our findings highlight the importance of adequate protection for humpback whales of the Kitimat Fjord System; the value of protected areas in general for any migratory species with strong seasonal site fidelity [126,189]; and the importance of long-term monitoring for the early detection and improved interpretation of population-level responses to changing marine ecosystems [108,126,170].

## Acknowledgements

The authors thank the Save Our Seas Foundation, Willow Grove Foundation, Donner Canadian Foundation, Tides Canada, LUSH Charity Pot, The Zumwalt Family, Julie Walters, Fisheries and Oceans Canada. Special acknowledgement is due to the Gitga’at First Nation of British Columbia, without their support this project would not have been possible.

## REFERENCES

1. Morris DW. Ecological Scale and Habitat Use. Ecology. 1987;68: 362–369.

2. Hutchinson GE. Concluding remarks. Cold Spring Harb Symp Quant Biol. 1957;22: 415–427.

3. Leibold MA. The Niche Concept Revisited: Mechanistic Models and Community Context. Ecology. 1995;76: 1371–1382.

4. Finke DL, Snyder WE. Niche partitioning increases resource exploitation by diverse communities. Science. 2008;321: 1488–1490.

5. Bolnick DI, Svanbäck R, Fordyce JA, Yang LH, Davis JM, Hulsey CD, et al. The ecology of individuals: incidence and implications of individual specialization. Am Nat. 2003;161: 1–28.

6. Emlen ST, Oring LW. Ecology, sexual selection, and the evolution of mating systems. Science. 1977;197: 215–223.

7. Gould SJ, Lewontin RC. The spandrels of San Marco and the Panglossian paradigm: a critique of the adaptationist programme. Proc R Soc Lond B Biol Sci. 1979;205: 581–598.

8. Gerber LR. Including behavioral data in demographic models improves estimates of population viability. Front Ecol Environ. 2006;4: 419–427.

9. Cañadas A, Hammond PS. Abundance and habitat preferences of the short-beaked common dolphin Delphinus delphis in the southwestern Mediterranean: implications for conservation. Endanger Species Res. 2008.

10. Musick JA, Lutz PL, Wyneken J. The biology of sea turtles. CRC Press; 2003.

11. Trillmich F, Trillmich KGK. The mating systems of pinnipeds and marine iguanas: convergent evolution of polygyny. Biol J Linn Soc Lond. 1984;21: 209–216.

12. Le Boeuf BJ. Pinniped mating systems on land, ice and in the water: Emphasis on the Phocidae. In: Renouf D, editor. The Behaviour of Pinnipeds. Dordrecht: Springer Netherlands; 1991. pp. 45–65.

13. Schreiber EA, Burger J. Biology of marine birds. CRC press; 2001.

14. Moehlman PD. Feral asses (Equus africanus): intraspecific variation in social organization in arid and mesic habitats. Appl Anim Behav Sci. 1998;60: 171–195.

15. Weir JS, Duprey NMT, Würsig B. Dusky dolphin (Lagenorhynchus obscurus) subgroup distribution: are shallow waters a refuge for nursery groups? Canadian Journal of. 2008.

16. Whitehead H, Rendell L. Movements, habitat use and feeding success of cultural clans of South Pacific sperm whales. J Anim Ecol. 2004; 190–196.

17. Clapham PJ. Humpback whale: Megaptera novaeangliae. In: Perrin WF, Würsig B, Thewissen JGM, editors. Encyclopedia of marine mammals (2nd ed). Academic Press; 2009. pp. 368–371.

18. Herman LM, Forestell PH, Antinoja RC. The 1976/77 migration of humpback whales into Hawaiian waters: composite description. 1980.

19. Glockner-Ferrari DA, Ferrari MJ. Individual Identification, behavior, reproduction, and distribution of humpback whales, Megaptera novaeangliae, in Hawaii. US Marine Mammal Commission; 1985.

20. Salden DR. Humpback whale encounter rates offshore of Maui, Hawaii. J Wildl Manage. 1988; 301– 304.

21. Glockner-Ferrari DA, Ferrari MJ. Reproduction in the humpback whale (Megaptera novaeangliae) in Hawaiian waters, 1975--1988: the life history, reproductive rates and behavior of known individuals identified through surface and underwater photography. Reports of the International Whaling Commission (Special Issue). 1990;12: 161–169.

22. Smultea MA. Segregation by humpback whale (Megaptera novaeangliae) cows with a calf in coastal habitat near the island of Hawaii. Can J Zool. 1994;72: 805–811.

23. Martins CCA, Morete ME, Coitinho MHE, Freitas AC, Secchi ER, Kinas PG. Aspects of habitat use patterns of humpback whales in the Abrolhos Bank, Brazil, breeding ground. 2001. Available: http://repositorio.furg.br/handle/1/1106

24. Frankel AS, Clark CW. ATOC and other factors affecting the distribution and abundance of humpback whales (Megaptera novaeangliae) off the north shore of Kauai. Mar Mamm Sci. 2002;18: 644– 662.

25. Ersts PJ, Rosenbaum HC. Habitat preference reflects social organization of humpback whales (Megaptera novaeangliae) on a wintering ground. J Zool. 2003;260: 337–345.

26. Gabriele CM, Rickards SH, Yin SE, Frankel AS. Trends in Relative Distribution, Abundance and Population Composition of Humpback Whales, Megaptera novaeangliae, in Kawaihae Bay, Hawai’i 1988-2003. 2003.

27. Félix F, Botero-Acosta N. Distribution and behaviour of humpback whale mother--calf pairs during the breeding season off Ecuador. Mar Ecol Prog Ser. 2011;426: 277–287.

28. Cartwright R, Gillespie B, Labonte K, Mangold T, Venema A, Eden K, et al. Between a rock and a hard place: habitat selection in female-calf humpback whale (Megaptera novaeangliae) Pairs on the Hawaiian breeding grounds. PLoS One. 2012;7: e38004.

29. Pack AA, Herman LM, Craig AS, Spitz SS, Waterman JO, Herman EYK, et al. Habitat preferences by individual humpback whale mothers in the Hawaiian breeding grounds vary with the age and size of their calves. Anim Behav. 2017;133: 131–144.

30. Würsig B. Delphinid foraging strategies. In: Schusterman RJ, Thomas JA, Wood FG, editors. Dolphin cognition and behavior: a comparative approach. Hillsdale, NJ: Lawrence Erlbaum Associates.; 987. pp. 347–359.

31. Whitehead H, Ford JKB. Consequences of culturally-driven ecological specialization: Killer whales and beyond. J Theor Biol. 2018;456: 279–294.

32. Ford JKB. Killer Whales: Behavior, Social Organization, and Ecology of the Oceans’ Apex Predators. Ethology and Behavioral Ecology of Odontocetes. Springer; 2019. pp. 239–259.

33. Whitehead H. Analysing animal social structure. Anim Behav. 1997;53: 1053–1067.

34. Jurasz CM, Jurasz VP. Feeding modes of the humpback whale, Megaptera novaeangliae, in Southeast Alaska. 1979.

35. Mastick N. The Effect of Group Size on Individual Roles and the Potential for Cooperation in Group Bubble-net Feeding Humpback Whales (Megaptera novaeangliae). 2016.

36. Foote AD, Newton J, Ávila-Arcos MC, Kampmann M-L, Samaniego JA, Post K, et al. Tracking niche variation over millennial timescales in sympatric killer whale lineages. Proc Biol Sci. 2013;280: 20131481.

37. Reed JM, Dobson AP.Behavioural constraints and conservation biology: Conspecific attraction and recruitment. Trends Ecol Evol. 1993;8: 253–256.

38. Whitehead H, Rendell L, Osborne RW, Würsig B. Culture and conservation of non-humans with reference to whales and dolphins: review and new directions. Biol Conserv. 2004;120: 427–437.

39. Grassi C. Variability in habitat, diet, and social structure of Hapalemur griseus in Ranomafana National Park, Madagascar. Am J Phys Anthropol. 2006;131: 50–63.

40. Pirotta E, Brotons JM, Cerdà M, Bakkers S, Rendell LE. Multi-scale analysis reveals changing distribution patterns and the influence of social structure on the habitat use of an endangered marine predator, the sperm whale Physeter macrocephalus in the Western Mediterranean Sea. Deep Sea Res Part I. 2020;155: 103169.

41. Boyd R, Richerson PJ. Why culture is common, but cultural evolution is rare. Proceedings-British Academy. Oxford University Press Inc.; 1996. pp. 77–94.

42. Laland KN, Hoppitt W. Do animals have culture? Evolutionary Anthropology: Issues, News, and Reviews. 2003;12: 150–159.

43. Whitehead H. Sperm whales: social evolution in the ocean. University of Chicago press; 2003.

44. Cantor M, Whitehead H. The interplay between social networks and culture: theoretically and among whales and dolphins. Philos Trans R Soc Lond B Biol Sci. 2013;368: 20120340.

45. Eguiguren A, Pirotta E, Cantor M, Rendell L, Whitehead H. Habitat use of culturally distinct Galápagos sperm whale Physeter macrocephalus clans. Mar Ecol Prog Ser. 2019;609: 257–270.

46. Geist V. Mountain sheep. A study in behavior and evolution. University of Chicago Press.; 1971.

47. Estes JA, Riedman ML, Staedler MM, Tinker MT, Lyon BE. Individual variation in prey selection by sea otters: patterns, causes and implications. J Anim Ecol. 2003; 44–155.

48. Boesch C, Marchesi P, Marchesi N, Fruth B, Joulian F. Is nut cracking in wild chimpanzees a cultural behaviour? J Hum Evol. 1994;26: 325–338.

49. van Schaik CP, Ancrenaz M, Borgen G, Galdikas B, Knott CD, Singleton I, et al. Orangutan cultures and the evolution of material culture. Science. 2003;299: 102–105.

50. Ottoni EB, Izar P. Capuchin monkey tool use: Overview and implications. Evolutionary Anthropology: Issues, News, and Reviews. 2008;17: 171–178.

51. Slagsvold T, Wiebe KL. Social learning in birds and its role in shaping a foraging niche. Philos Trans R Soc Lond B Biol Sci. 2011;366: 969–977.

52. Keen SC, Cole EF, Sheehan MJ, Sheldon BC. Social learning of acoustic anti-predator cues occurs between wild bird species. Proc Biol Sci. 2020;287: 20192513.

53. Rendell L, Whitehead H. Spatial and temporal variation in sperm whale coda vocalizations: stable usage and local dialects. Anim Behav. 2005;70: 191–198.

54. Marcoux M. Vocalizations, diet and fitness among acoustic clans of sperm whales (Physeter macrocephalus). M.Sc., Dalhousie University (Canada). 2005.

55. Marcoux M, Whitehead H, Rendell L. Sperm whale feeding variation by location, year, social group and clan: evidence from stable isotopes. Mar Ecol Prog Ser. 2007;333: 309–314.

56. Cantor M, Whitehead H. How does social behavior differ among sperm whale clans?Mar Mamm Sci. 2015;31: 1275–1290.

57. Hauser DDW, Logsdon MG, Holmes EE, VanBlaricom GR, Osborne RW. Summer distribution patterns of southern resident killer whales Orcinus orca: core areas and spatial segregation of social groups. Mar Ecol Prog Ser. 2007;351: 301–310.

58. Chilvers LB, Corkeron PJ. Trawling and bottlenose dolphins’ social structure. Proceedings of the Royal Society of London Series B: Biological Sciences. 2001;268: 1901–1905.

59. Markowitz TM, Harlin AD, Würsig B, McFadden CJ. Dusky dolphin foraging habitat: overlap with aquaculture in New Zealand. Aquat Conserv. 2004;14: 133–149.

60. Mann J, Stanton MA, Patterson EM, Bienenstock EJ, Singh LO. Social networks reveal cultural behaviour in tool-using dolphins. Nat Commun. 2012;3: 1–8.

61. Clapham PJ, Mayo CA. Reproduction and recruitment of individually identified humpback whales, Megaptera novaeangliae, observed in Massachusetts Bay, 1979–1985. Can J Zool. 1987;65: 2853–2863.

62. Valenzuela LO, Sironi M, Rowntree VJ, Seger J. Isotopic and genetic evidence for culturally inherited site fidelity to feeding grounds in southern right whales (Eubalaena australis). Mol Ecol. 2009;18: 782– 791.

63. Baker CS, Steel D, Calambokidis J, Falcone E, González-Peral U, Barlow J, et al. Strong maternal fidelity and natal philopatry shape genetic structure in North Pacific humpback whales. Mar Ecol Prog Ser. 2013;494: 291–306.

64. Weitkamp LA, Wissmar RC. Gray whale foraging on ghost shrimp (Callianassa californiensis) in littoral sand flats of Puget Sound, USA. J Zool. 1992.

65. Allen J, Weinrich M, Hoppitt W, Rendell L. Network-based diffusion analysis reveals cultural transmission of lobtail feeding in humpback whales. Science. 2013;340: 485–488.

66. McMillan CJ, Towers JR, Hildering J. The innovation and diffusion of “trap-feeding,” a novel humpback whale foraging strategy. Mar Mamm Sci. 2019;35: 779–796.

67. Dawbin WH. The seasonal migratory cycle of humpback whales. Whales, dolphins and porpoises. 1966; 145–170.

68. Weinrich MT, Rosenbaum H, Scott Baker C, Blackmer AL, Whitehead H. The influence of maternal lineages on social affiliations among humpback whales (Megaptera novaeangliae) on their feeding grounds in the southern Gulf of Maine. J Hered. 2006;97: 226–234.

69. Connor RC. Group living in whales and dolphins. In: Mann J, Connor RC, Tyack PL, Whitehead H, editors. Cetacean societies: field studies of dolphins and whales. Chicago: University of Chicago Press; 2000. pp. 199–218.

70. Clapham PJ. The social and reproductive biology of Humpback Whales: an ecological perspective. Mamm Rev. 1996;26: 27–49.

71. Whitehead H. Structure and stability of humpback whale groups off Newfoundland. Can J Zool. 1983.

72. Weinrich MT. Long term stability in grouping patterns of humpback whales (Megaptera novaeangliae) in the southern Gulf of Maine. Can J Zool. 1991;69: 3012–3019.

73. Weinrich MT, Kuhlberg AE. Short-term association patterns of humpback whale (Megaptera novaeangliae) groups on their feeding grounds in the southern Gulf of Maine. Can J Zool. 1991;69: 3005– 3011.

74. Baker CS. The population structure and social organization of humpback whales (Megaptera novaeangliae) in the central and eastern North Pacific. University of Hawaii. 1985.

75. Mobley JR Jr, Herman LM. Transience of social affiliations among humpback whales (Megaptera novaeangliae) on the Hawaiian wintering grounds. Can J Zool. 1985;63: 762–772.

76. Mattila DK, Clapham PJ, Vásquez O, Bowman RS. Occurrence, population composition, and habitat use of humpback whales in Samana Bay, Dominican Republic. Can J Zool. 1994;72: 1898–1907.

77. Ramp C, Hagen W, Palsbøll P, Bérubé M, Sears R. Age-related multi-year associations in female humpback whales (Megaptera novaeangliae). Behav Ecol Sociobiol. 2010;64: 1563–1576.

78. Lubansky TM. E pluribus unum: what individual whales can tell us about enigmatic species distribution and social organization. New Jersey Institute of Technology. 2015.

79. Brakes P, Dall SRX, Aplin LM, Bearhop S, Carroll EL, Ciucci P, et al. Animal cultures matter for conservation. Science. 2019;363: 1032–1034.

80. Nichol LM, Abernethy R, Flostrand L, Lee TS, Ford J, of Fisheries D. Information relevant to the identification of critical habitats of North Pacific humpback whales(Megaptera novaeangliae) in British Columbia. DFO, Ottawa, ON(Canada); 2010.

81. Ashe E, Wray J, Picard CR, Williams R. Abundance and survival of Pacific humpback whales in a proposed critical habitat area. PLoS One. 2013;8: e75228.

82. Calambokidis J, Falcone EA, Quinn TJ, Burdin AM, Clapham PJ, Ford JKB, et al. SPLASH: Structure of Populations, Levels of Abundance and Status of Humpback Whales in the North Pacific. U.S. Dept of Commerce; 2008 May. Available: http://www.cascadiaresearch.org/files/publications/SPLASH-contract-Report-May08.pdf

83. Keen EM. bangarang. 2016. Available: https://github.com/ericmkeen/bangarang

84. Keen EM, Wray J, Meuter H, Thompson K-L, Barlow JP, Picard CR. “Whale wave”: shifting strategies structure the complex use of critical fjord habitat by humpbacks. Mar Ecol Prog Ser. 2017;567: 211–233.

85. Keen EM, Wray J, Pilkington JF, Thompson KL, Picard CR. Distinct habitat use strategies of sympatric rorqual whales within a fjord system. Mar Environ Res. 2018;140: 180–189.

86. R Core Team. R: A language and environment for statistical computing. 2020. Available: https://www.R-project.org/

87. Baker CS. Seasonal contrasts in the social-behavior of the North Pacific humpback whale. Pacific Science. Univ Hawaii Press 2840 Kolowalu St, Honolulu, HI 96822; 1984. pp. 356–356.

88. Baker CS, Herman LM. Aggressive behavior between humpback whales (Megaptera novaeangliae) wintering in Hawaiian waters. Can J Zool. 1984.

89. Mattila DK, Clapham PJ. Humpback whales, Megaptera novaeangliae, and other cetaceans on Virgin Bank and in the northern Leeward Islands, 1985 and 1986. Can J Zool. 1989;67: 2201–2211.

90. Clapham PJ. Social organization of humpback whales on a North Atlantic feeding ground. Symposia of the Zoological Society of London. 1993. pp. 131–145.

91. White GC, Garrott RA. Analysis of Wildlife Radio-tracking Data. California: Academic Press: San Diego; 1990.

92. Whitehead H. Investigating structure and temporal scale in social organizations using identified individuals. Behav Ecol. 1995;6: 199–208.

93. Perryman RJY, Venables SK, Tapilatu RF, Marshall AD, Brown C, Franks DW. Social preferences and network structure in a population of reef manta rays. Behav Ecol Sociobiol. 2019;73: 114.

94. Whitehead H. Analyzing animal societies: quantitative methods for vertebrate social analysis. University of Chicago Press; 2008.

95. Farine DR. A guide to null models for animal social network analysis. Methods Ecol Evol. 2017;8: 1309–1320.

96. Whitehead H. SOCPROG: programs for analyzing social structure. 2019. Available: http://whitelab.biology.dal.ca/SOCPROG/Manual.pdf

97. Whitehead H. Analysis of animal movement using opportunistic individual identifications: application to sperm whales. Ecology. 2001;82: 1417–1432.

98. Tschopp A, Ferrari MA, Crespo EA, Coscarella MA. Development of a site fidelity index based on population capture-recapture data. PeerJ. 2018;6: e4782.

99. Acevedo J, Mora C, Aguayo-Lobo A. Sex-related site fidelity of humpback whales (Megaptera novaeangliae) to the Fueguian Archipelago feeding area, Chile. Mar Mamm Sci. 2014;30: 433–444.

100. Cairns SJ, Schwager SJ. A comparison of association indices. Anim Behav. 1987;35: 1454–1469.

101. Hoppitt WJE, Farine DR. Association indices for quantifying social relationships: how to deal with missing observations of individuals or groups. Anim Behav. 2018;136: 227–238.

102. Weko CW. Isolating bias in association indices. Anim Behav. 2018;139: 147–159.

103. Farine DR. Animal social network inference and permutations for ecologists in R using asnipe. O’Hara RB, editor. Methods Ecol Evol. 2013;4: 1187–1194.

104. Whitehead H, James R. Generalized affiliation indices extract affiliations from social network data. Methods Ecol Evol. 2015;6: 836–844.

105. Godde S, Humbert L, Côté SD, Réale D, Whitehead H. Correcting for the impact of gregariousness in social network analyses. Anim Behav. 2013;85: 553–558.

106. Csardi G, Nepusz T. The igraph software package for complex network research. InterJournal, Complex Systems 1695. 2006. Available: http://igraph.org

107. Blondel VD, Guillaume J-L, Lambiotte R, Lefebvre E. Fast unfolding of communities in large networks. J Stat Mech: Theory Exp. 2008;2008: P10008.

108. Wray J, Keen EM. Calving rate decline in humpback whales (Megaptera novaeangliae) of northern British Columbia, Canada. Mar Mamm Sci. 2020;36: 709–720.

109. Rand WM. Objective criteria for the evaluation of clustering methods. J Am Stat Assoc. 1971;66: 846–850.

110. Hubert L, Arabie P. Comparing partitions. J Classification. 1985;2: 193–218.

111. Csárdi G, Nepusz T, Airoldi EM. Statistical network analysis with igraph. New York, NY: Springer; 2016.

112. Farine DR. Assortnet. 2016. Available: https://cran.r-project.org/web/packages/assortnet/assortnet.pdf

113. Farine DR. Measuring phenotypic assortment in animal social networks: weighted associations are more robust than binary edges. Anim Behav. 2014;89: 141–153.

114. Opsahl T, Agneessens F, Skvoretz J. Node centrality in weighted networks: Generalizing degree and shortest paths. Soc Networks. 2010;32: 245–251.

115. Derville S, Torres LG, Garrigue C. Social segregation of humpback whales in contrasted coastal and oceanic breeding habitats. J Mammal. 2018;99: 41–54.

116. Calambokidis J, Steiger GH, Straley JM, Herman LM, Cerchio S, Salden DR, et al. Movements and population structure of humpback whales in the North Pacific. Mar Mamm Sci. 2001;17: 769–794.

117. Taylor BL. The Reliability of Using Population Viability Analysis for Risk Classification of Species. Conserv Biol. 1995;9: 551–558.

118. Palsbøll PJ, Clapham PJ, Mattila DK, Larsen F, Sears R, Siegismund HR, et al. Distribution of mtDNA haplotypes in North Atlantic humpback whales: the influence of behaviour on population structure. Mar Ecol Prog Ser. 1995;116: 1–10.

119. Barendse J, Best PB, Carvalho I, Pomilla C. Mother knows best: occurrence and associations of resighted humpback whales suggest maternally derived fidelity to a Southern Hemisphere coastal feeding ground. PLoS One. 2013;8: e81238.

120. Stevick PT, Allen J, Clapham PJ, Katona SK, Larsen F, Lien J, et al. Population spatial structuring on the feeding grounds in North Atlantic humpback whales (Megaptera novaeangliae). J Zool. 2006;270: 244–255.

121. Barlow J, Calambokidis J, Falcone EA, Baker CS, Burdin AM, Clapham PJ, et al. Humpback whale abundance in the North Pacific estimated by photographic capture-recapture with bias correction from simulation studies. Mar Mamm Sci. 2011;27: 793–818.

122. Ford JKB. Marine Mammals of British Columbia. Royal BC Museum Victoria, British Columbia; 2014.

123. Wade PR, Quinn TJ, Barlow J, Baker CS, Burdin AM, Calambokidis J, et al. Estimates of abundance and migratory destination for North Pacific humpback whales in both summer feeding areas and winter mating and calving areas. International Whaling Commission Report SC/66b/IA/21. 2016.

124. Carretta JV, Oleson EM, Weller DW, Lang AR, Forney KA, Baker JD, et al. US Pacific marine mammal stock assessments, 2013. 2014.

125. Baker CS, Herman LM, Perry A, Lawton WS, Straley JM, Wolman AA, et al. Migratory movement and population structure of humpback whales (Megaptera novaeangliae) in the central and eastern North Pacific. Mar Ecol Prog Ser. 1986;31: 105–119.

126. Gabriele CM, Neilson JL, Straley JM, Baker CS, Cedarleaf JA, Saracco JF. Natural history, population dynamics, and habitat use of humpback whales over 30 years on an Alaska feeding ground. Ecosphere. 2017;8. doi:10.1002/ecs2.1641

127. Riley HE. Humpback whale (Megaptera novaeangliae) numbers and distribution on their summer feeding grounds of the Eastern Aleutian Islands. University of Alaska Fairbanks. 2010.

128. Witteveen BH, Wynne KM. Site fidelity and movement of humpback whales (Megaptera novaeangliae) in the western Gulf of Alaska as revealed by photo-identification. Can J Zool. 2017;95: 169–175.

129. Hucke-Gaete R, Haro D, Torres-Florez JP, Montecinos Y, Viddi F, Bedriñana-Romano L, et al. A historical feeding ground for humpback whales in the eastern South Pacific revisited: the case of northern Patagonia, Chile. Aquat Conserv. 2013;23: 858–867.

130. Clapham PJ, Baraff LS, Carlson CA, Christian MA, Mattila DK, Mayo CA, et al. Seasonal occurrence and annual return of humpback whales, Megaptera novaeangliae, in the southern Gulf of Maine. Can J Zool. 1993;71: 440–443.

131. Herman LM, Pack AA, Rose K, Craig A. Resightings of humpback whales in Hawaiian waters over spans of 10–32 years: Site fidelity, sex ratios, calving rates, female demographics, and the dynamics of social and behavioral roles of individuals. Marine Mammal Science. 2011;27: 736–768.

132. Wedekin LL, Neves MC, Marcondes MCC, Baracho C, Rossi-Santos MR, Engel MH, et al. Site fidelity and movements of humpback whales (Megaptera novaeangliae) on the Brazilian breeding ground, southwestern Atlantic. Mar Mamm Sci. 2010;26: 787–802.

133. Baracho-Neto CG, Neto ES, Rossi-Santos MR, Wedekin LL, Neves MC, Lima F, et al. Site fidelity and residence times of humpback whales (Megaptera novaeangliae) on the Brazilian coast. J Mar Biol Assoc U K. 2012;92: 1783.

134. Craig AS, Herman LM. Sex differences in site fidelity and migration of humpback whales (Megaptera novaeangliae) to the Hawaiian Islands. Can J Zool. 1997;75: 1923–1933.

135. Scheidat M, Castro C, Denkinger J, González J, Adelung D. A breeding area for humpback whales (Megaptera novaeangliae) off Ecuador. J Cetacean Res Manag. 2000;2: 165–172.

136. Baker CS, Herman LM. Migration and local movement of humpback whales (Megaptera novaeangliae) through Hawaiian waters. Can J Zool. 1981;59: 460–469.

137. Acebes JMV, Darling JD, Yamaguchi M. Status and distribution of humpback whales (Megaptera novaeangliae) in northern Luzon, Philippines. J Cetacean Res Manage Special Issue. 2007;9: 37–43.

138. Keen EM. Aggregative and feeding thresholds of sympatric rorqual whales within a fjord system. Ecosphere. 2017;8: e01702.

139. Hain JHW, Carter GR, Kraus SD, Mayo CA, Winn HE. Megaptera Novaeangliae, in the Western North Atlantic. Fish Bull. 1982;80: 259.

140. D’Vincent CG, Nilson RM, Hanna RE. Vocalization and coordinated feeding behavior of the humpback whale in southeastern Alaska. Sci Rep Whales Res Inst. 1985;36: 41–47.

141. Clapham PJ. The Humpback Whale: Seasonal Feeding and Breeding in a Baleen Whale. In: Mann J, editor. Field Studies of Dolphins and Whales. University of Chicago Press; 2000. pp. 173–196.

142. Wiley D, Ware C, Bocconcelli A, Cholewiak D, Friedlaender A, Thompson M, et al. Underwater components of humpback whale bubble-net feeding behaviour. Behaviour. 2011;148: 575–602.

143. Sharpe FA, Dill LM. The behavior of Pacific herring schools in response to artificial humpback whale bubbles. Can J Zool. 1997;75: 725–730.

144. Perry A, Baker CS, Herman LM. Population characteristics of individually identified humpback whales in the central and eastern North Pacific: a summary and critique. Repts intl Whaling Commn Special Issue. 1990; 307–317.

145. Sharpe FA. Social foraging of the southeast Alaskan humpback whale, Megaptera novaeangliae. Ph.D., Simon Fraser University. 2001.

146. Tyack P, Whitehead H. Male Competition in Large Groups of Wintering Humpback Whales. Behaviour. 1983;83: 132–154.

147. Palsbøll PJ, Clapham PJ, Mattila DK, Vasquez O. Composition and Dynamics of Humpback Whale Competitive Groups in the West Indies. Behaviour. 1992;122: 182–194.

148. Perry A, Baker CS, Herman LM. Population characteristics of individually identified humpback whales in the central and eastern North Pacific: a summary and critique. Rep Int Whal Commn (Special Issue). 1990; 307–317.

149. Krieger KJ, Wing BL. Hydroacoustic surveys and identification of humpback whale forage in Glacier Bay, Stephens Passage, and Frederick Sound, southeastern Alaska, summer 1983. NOAA; 1984.

150. Baker CS, Herman LM, Perry A, Lawton WS, Straley JM, Straley JH. Population characteristics and migration of summer and late-season humpback whales (Megaptera novaeangliae) in southeastern Alaska. Mar Mamm Sci. 1985;1: 304–323.

151. Perry A, Baker CS, Herman LM. The natural history of humpback whales in Glacier Bay, Alaska. National Park Service, Alaska Regional Office, Anchorage; 1985.

152. Cartwright R, Venema A, Hernandez V, Wyels C, Cesere J, Cesere D. Fluctuating reproductive rates in Hawaii’s humpback whales, Megaptera novaeangliae, reflect recent climate anomalies in the North Pacific. R Soc Open Sci. 2019;6: 181463.

153. Bagulayan A, Bartlett-Roa JN, Carter AL, Inman BG, Keen EM, Orenstein EC, et al. Journey to the center of the gyre: The fate of the Tohoku Tsunami debris field. Oceanography. 2012;25: 200–207.

154. Bond NA, Cronin MF, Freeland H, Mantua N. Causes and impacts of the 2014 warm anomaly in the NE Pacific. Geophys Res Lett. 2015;42: 3414–3420.

155. Dewey R. The Blob Blog - Warm Northeast Pacific Ocean Conditions Continue (2016). Feb 2016 [cited 26 Nov 2020]. Available: https://www.oceannetworks.ca/blob-blog-warm-northeast-pacific-ocean-conditions-continue-2016

156. Jones T, Parrish JK, Peterson WT, Bjorkstedt EP, Bond NA, Ballance LT, et al. Massive Mortality of a Planktivorous Seabird in Response to a Marine Heatwave. Geophys Res Lett. 2018;45: 3193–3202.

157. Walsh JE, Thoman RL, Bhatt US, Bieniek PA, Brettschneider B, Brubaker M, et al. The high latitude marine heat wave of 2016 and its impacts on Alaska. Bulletin of the American Meteorological Society. 2018;99: 39–43.

158. Tanner CJ, Jackson AL. The combination of social and personal contexts affects dominance hierarchy development in shore crabs, Carcinus maenas. Anim Behav. 2011;82: 1185–1192.

159. Henzi SP, Lusseau D, Weingrill T, van Schaik CP, Barrett L. Cyclicity in the structure of female baboon social networks. Behav Ecol Sociobiol. 2009;63: 1015–1021.

160. Ramirez-Sanchez S, Pinkerton E. The impact of resource scarcity on bonding and bridging social capital: the case of fishers’ information-sharing networks in Loreto, BCS, Mexico. Ecol Soc. 2009;14.

161. Beck S, Kuningas S, Esteban R, Foote AD. The influence of ecology on sociality in the killer whale (Orcinus orca). Behav Ecol. 2012;23: 246–253.

162. Foster EA, Franks DW, Morrell LJ, Balcomb KC, Parsons KM, van Ginneken A, et al. Social network correlates of food availability in an endangered population of killer whales, Orcinus orca. Anim Behav. 2012;83: 731–736.

163. Williams R, Lusseau D, Hammond PS. The role of social aggregations and protected areas in killer whale conservation: The mixed blessing of critical habitat. Biol Conserv. 2009;142: 709–719.

164. Durant SM. Competition refuges and coexistence: an example from Serengeti carnivores. J Anim Ecol. 1998;67: 370–386.

165. Sutherland WJ. The importance of behavioural studies in conservation biology. Anim Behav. 1998;56: 801–809.

166. McComb K, Moss C, Durant SM, Baker L, Sayialel S. Matriarchs as repositories of social knowledge in African elephants. Science. 2001;292: 491–494.

167. Baird RW, Gorgone AM, McSweeney DJ, Webster DL, Salden DR, Deakos MH, et al. False killer whales (Pseudorca crassidens) around the main Hawaiian Islands: Long-term site fidelity, inter-island movements, and association patterns. Mar Mamm Sci. 2008;24: 591–612.

168. Baird RW, Hanson MB, Schorr GS, Webster DL. Range and primary habitats of Hawaiian insular false killer whales: informing determination of critical habitat. Endanger Species Res. 2012.

169. Mcdonald RI, Kareiva P, Forman RTT. The implications of current and future urbanization for global protected areas and biodiversity conservation. Biol Conserv. 2008;141: 1695–1703.

170. Hoegh-Guldberg O, Bruno JF. The impact of climate change on the world’s marine ecosystems. Science. 2010;328: 1523–1528.

171. Macdonald RW, Bornhold BD, Webster I. The Kitimat fjord system: An introduction. in Proceedings of a workshop on the Kitimat marine environment. Canadian Technical Report of Hydrography and Ocean Sciences No 18 Institute of Ocean Sciences, Department of Fisheries and Oceans. 1983;18: 2–13.

172. Hoekstra JM, Fagan WF, Bradley JE. A critical role for critical habitat in the recovery plan process? Not yet. Ecol Appl. 2002;12: 701–707.

173. Clark JA, Hoekstra JM, Boersma PD, Kareiva P. Improving US Endangered Species Act recovery plans: key findings and recommendations of the SCB recovery plan project. Conserv Biol. 2002;16: 1510–1519.

174. Taylor MFJ, Suckling KF, Rachlinski JJ. The effectiveness of the Endangered Species Act: a quantitative analysis. Bioscience. 2005;55: 360–367.

175. Reed JM, Akcakaya HR, Burgman M, Bender D, Beissinger SR, Scott JM. Critical habitat. In: Goble D, Scott JM, Davis FW, editors. The Endangered Species Act at Thirty: Conserving biodiversity in human-dominated landscapes Evolution of at-risk species protection. Washington, D.C.: Island Press; 2006. pp. 164–177.

176. Schwartz MW. The performance of the endangered species act. Annu Rev Ecol Evol Syst. 2008;39.

177. Gibbs KE, Currie DJ. Protecting endangered species: do the main legislative tools work? PLoS One. 2012;7: e35730.

178. Camaclang AE, Maron M, Martin TG, Possingham HP. Current practices in the identification of critical habitat for threatened species. Conserv Biol. 2015;29: 482–492.

179. District of Kitimat Development Services (DKDS). The Private International Port of Kitimat. Kitimat: A Port City on the Move, 1-20. Yumpu.com; 2015 [cited 26 Nov 2020]. Available: https://www.yumpu.com/en/document/read/34172480/the-private-international-port-of-kitimat

180. Angevine G, Oviedo V. Laying the Groundwork for BC LNG Exports to Asia. Studies in Energy Policy, Fraser Institute. 2012.

181. Meisner T. TERMPOL Review Process on LNG CANADA Project. (Report No. TP 15287E, Catalogue No. T29-126/2015E). Transport Canada; 2015. Available: http://publications.gc.ca/collections/collection_2016/tc/T29-126-2015-eng.pdf

182. Erbe C, Peel D, Redfern J, Smith JN. Impacts of shipping on marine fauna. Frontiers in Marine Science; 2020.

183. Hatch L, Clark C, Merrick R, Van Parijs S, Ponirakis D, Schwehr K, et al. Characterizing the relative contributions of large vessels to total ocean noise fields: a case study using the Gerry E. Studds Stellwagen Bank National Marine Sanctuary. Environmental Management. 2008;42: 735–752.

184. Erbe C. Effects of underwater noise on marine mammals. Adv Exp Med Biol. 2012;730: 17–22.

185. Payne R, Webb D. Orientation by means of long range acoustic signaling in baleen whales. Ann N Y Acad Sci. 1971;188: 110–141.

186. Roman J, Estes JA, Morissette L, Smith C, Costa D, McCarthy J, et al. Whales as marine ecosystem engineers. Front Ecol Environ. 2014;12: 377–385.

187. Francis JM. Nature conservation and the precautionary principle. Environ Values. 1996; 257–264.

188. Gerrodette T, Dayton PK, Macinko S, Fogarty MJ. Precautionary management of marine fisheries: moving beyond burden of proof. Bull Mar Sci. 2002;70: 657–668.

189. Bräger S, Dawson SM, Slooten E, Smith S, Stone GS, Yoshinaga A. Site fidelity and along-shore range in Hector’s dolphin, an endangered marine dolphin from New Zealand. Biol Conserv. 2002;108: 281–287.

